# Complement-Activating Multimeric Immunotherapeutic Complexes for HER2-breast cancer immunotherapy

**DOI:** 10.1101/2024.02.02.578619

**Authors:** Carole Seguin-Devaux, Bianca Brandus, Jean-Marc Plesseria, Gilles Iserentant, Jean-Yves Servais, Georgia Kanli, Iris Behrmann, Jacques Zimmer, Jacques H M Cohen, Xavier Dervillez

## Abstract

**Background:** Directing selective complement activation towards tumor cells is an attractive strategy to promote their elimination. We have generated Complement-activating Multimeric immunotherapeutic compleXes (CoMiX) that selectively stimulate the alternative pathway using Factor H Related protein 4 (FHR4) or the classical complement pathways using triple Fc dimers on HER2-expressing tumor cells.

**Methods:** We used the C4bp C-terminal-α-/β-chain multimerising scaffolds to generate CoMiX-FHR4 and CoMiX-Fc with 2 different V_H_H anti-HER2, V_H_H(T) and V_H_H(P), recognising trastuzumab-or pertuzumab-competing HER2 epitopes, respectively: FHR4/V_H_H(T), FHR4/V_H_H(P), V_H_H(T)/Fc, V_H_H(P)/Fc. The different CoMiX were compared *in vitro* for C3b and C5b9 depositions, complement-dependent cytotoxicity, and their ability to activate NK cells and phagocytosis by macrophages using one-way ANOVA and post-hoc Tukey’s tests. We further explored their therapeutic efficacy *in vivo* on human BT474 breast cancer xenografts established in NUDE mice, when used individually or in combination, as compared to trastuzumab or pertuzumab.

**Results:** FHR4/V_H_H(T) and FHR4/V_H_H(P) led to the highest C3b and C5b9 depositions and CDC, both individually and in combinations on BT474 tumor cells (p< 0.0001) surpassing the very low complement activating capacity of trastuzumab and pertuzumab. CoMiX-Fc showed NK cell activation and complement-mediated BT474 phagocytosis by M2 macrophages. In the xenograft model, CoMiX-FHR4 molecules reduced the tumor volume by a factor of 7.33 compared to the PBS control. V_H_H(T)/Fc had no effect on tumor growth, while V_H_H(P)/Fc led to a 2.75-times tumor volume reduction that was higher than pertuzumab (p< 0.01). Trastuzumab and its combination with pertuzumab remained the most potent regimen, alone or in combination, to completely inhibit tumor growth. CoMiX-FHR4, CoMiX-Fc and C3b deposition were visualized as soon as one hour after injection resulting in a massive homogeneous complement deposit 6 hours after injection. Interestingly, CoMiX-FHR4 significantly reduced the growth of trastuzumab-resistant cancer cells in contrast to trastuzumab and induced a large NK cell infiltration into the tumor.

**Conclusions:** CoMiX-FHR4 and CoMiX V_H_H(P)/Fc significantly inhibit tumor growth through complement activation, NK cells infiltration, and phagocytosis by macrophages. CoMiX-FHR4 proteins delay xenograft growth of BT474 cells resistant to trastuzumab and could thus be an attractive approach when resistance to antibody emerges.

**Key messages:** *What is already known on this topic:* Complement activation represents a substantial part of the overall biological activity of few therapeutic antibodies used in cancer immunotherapy. Factor H-related protein 4 can activate complement by serving as a platform for the assembly of alternative pathway C3 convertase by competing with factor H for C3b binding. We previously showed that multimeric recombinant proteins displaying the FHR4 complement effector moiety and a nanobody anti-HER2 targeting moiety selectively direct the activation of the complement alternative pathway on HER2-expressing tumor cells, leading to subsequent cell destruction through direct cell lysis or through the activation of host effector cells.

*What this study adds:* We used in the current work a novel complement-directed tumor cell distruction strategy *in vivo*. We showed that CoMiX-FHR4 and CoMiX-Fc (based on triple Fc dimers), targeting HER2-positive breast tumor cells, inhibit tumor growth in a model of BT474 xenograft in NUDE mice by stimulating complement activation, BT474 death, NK cell activation, and phagocytosis of tumor cells by macrophages. CoMiX-FHR4 remain efficient in xenografts of BT474 cells resistant to trastuzumab.

*How this study might affect research, practice or policy:* We demonstrate for the first time that directed complement activation on tumor cells is an alternative to therapeutic antibodies which is particularly promising when resistance to standard-of-care treatment occurs.

## Background

Recombinant monoclonal immunoglobulin gamma antibodies (IgGs) have become a major weapon in cancer immunotherapy. The classical complement pathway is activated when antibody–antigen complexes are formed on cell surfaces, followed by binding of C1q to the constant region of the antigen-bound antibody^1^. Few therapeutic antibodies (Abs), such as rituximab targeting CD20, efficiently recruit the complement protein C1q of the complement system which binds to the C-terminal half of the CH2 domain of Fc and leads to complement-dependent cytotoxicity (CDC)^2^. All three complement pathways, activated by antibodies or by structural elements on target cells or pathogens, converge at the level of C3 amplification loop, which is cleaved by C3 convertase to produce the anaphylatoxin C3a and the component C3b, followed by the sequential recruitment and activation of C5^2^. The assembly of the terminal complement proteins C5b–C9, the membrane attack complex (MAC), results in pore formation and subsequent lysis of the target cells. C3b degradation products on tumor targets (iC3b, C3dg and C3d^3^) are recognized by effector cells through CD11b and CD11c, leading to complement-dependent cell cytotoxicity (CDCC) or phagocytosis (CDCP), which allows the clearance of target cells^4^. The Fc region, in particular the CH2-CH3 domain interface, interacts also with the neonatal Fc receptor (FcRn) that has a quasi-ubiquitous expression pattern and modulates the transcytotic trafficking of IgGs. FcRn thus has a central role of a homeostatic regulator of the circulating levels of IgGs and dramatically increases the IgG half-life in circulation^5^. The Fab of therapeutic IgGs can also modulate the Tumor-associated Antigen (TAA)-linked downstream signaling pathway which may lead to a cytostatic effect and inhibition of tumor growth. Association of two therapeutic antibodies recognizing different epitopes of the same TAA can also lead to efficient complement activation, despite not activating complement when used individually. This is the case for trastuzumab that inhibits the human epidermal growth factor receptor 2 (HER2) and EGFR signalling pathways^6^.

Trastuzumab was the first humanized mAb approved for cancer treatment. Pertuzumab binds to a distinct epitope on HER2 but includes the same IgG1 Fc domain as trastuzumab. Both antibodies can trigger antibody-dependent cell-mediated cytotoxicity (ADCC), however, when combined, few additive effects were reported. Several critical factors determine the ability of an antibody to activate the classical complement pathway, including epitope recognition, size, target protein density, membrane fluidity, and the internalization kinetics of the targeted receptor following antibody-epitope interactions^7^. The ability to recruit C1q is also influenced by the orientation and geometry of the IgG following its binding to its epitope. Indeed, hexamerization of IgG antibodies following antigen fixation provides a platform for efficient C1q binding, facilitating significant complement activation^8^.

Nevertheless, despite the attractiveness of IgG therapies, many patients do not respond to current therapies, due to inherited or acquired resistance. There are still major medical needs to improve the efficacy of therapeutic mAbs. Modification of the effector functions using Fc bioengineering^9^ or antibody drug-conjugation combining anticancer drug such as trastuzumab-emtansine (T-DM1)^10^ are the main important approaches. The alternative pathway represents another critical component of the complement pathway that can be harnessed against tumor progression^11, 12^.

We have developed a novel anti-tumor complement-mediated strategy by exploiting the complement activating effects of FH-related protein 4 (FHR4) towards HER2-overexpressing cancer cells^13, 14^. FHR4 competes with factor H (FH) for C3b binding, resulting in the de-regulation of FH-mediated decay, thus leading to local activation of the complement amplification loop. We used the C4bp C-terminal α-chain multimerising scaffold (C4bpα) to generate <u>co</u>mplement-activating <u>m</u>ultimeric immunotherapeutic comple<u>x</u>es (CoMiX) that display multivalent FHR4 complement effector moieties and elicit C3b deposition, MAC formation and CDC by overcoming the complement inhibitory threshold on cancer cells ^13^. In the present work, we have extended the use of the C4bpα multimerizing scaffold to express CoMiX-Fc harbouring multimeric Fc that assemble as functional triple Fc-dimers. We used 2 different V_H_H anti-HER2, V_H_H(T) and V_H_H(P), recognising trastuzumab- or pertuzumab-competing HER2 epitopes, respectively, to generate 2 types of CoMiX-FHR4 molecules [FHR4/V_H_H(T) and FHR4/V_H_H(P)] and 2 types of CoMiX-Fc molecules [V_H_H(T)/Fc and V_H_H(P)/Fc]. CoMiX-FHR4 and CoMiX-Fc were first compared *in vitro* to trastuzumab and pertuzumab, used individually or in combination. The therapeutic efficacy of CoMiX-FHR4 and CoMiX-Fc was finally confirmed *in vivo* using xenografts established in NUDE mice from human HER2-expressing BT474 breast cancer cell lines that are sensitive or not to trastuzumab. Our study shows that the directed complement recruitment strategy is an efficient approach to inhibit tumor growth, and particularly promising in conditions of immunotherapeutic failure when tumors become resistant to standard treatment.

## Material and methods

### Transfection and clone selection

All CoMiX were generated from HEK293T cells transfected with four μg of recombinant plasmid DNA using Lipofectamine-3000 (Invitrogen, Thermo Fisher Scientific BVBA, Merelbeke, Belgium) following the manufacturer’s instructions. CoMiX-FHR4 were produced as previously described^13^ using double co-transfection whereas CoMiX-Fc, hinge deficient CoMiX-Fc (Δhinge) and multimeric V_H_H(T) control, lacking any FHR4 or Fc effector functions, were produced using a single transfection. After puromycin (Westburg, Leusden, The Netherlands) selection, individual puromycin-resistant cell clones were cultured. Protein expression in the supernatants were analysed using anti-Fc ELISA for CoMiX-Fc molecules, and symetrical anti-His ELISA for CoMiX-FHR4 molecules and V_H_H(T) control. 100 ng/well goat anti-human IgG Fc (Abcam, Cambridge, UK, #ab97221) or rabbit anti-His (Bethyl, ImTec Diagnostic NV, Antwerpen, Belgiu, #A190214A) pAbs were immobilized onto a NUNC MaxiSorp™ 96-well flat-bottom polystyrene plate overnight at 4°C. After blocking with 5% (w/v) bovine serum albumin in PBS, 1 µl of clone supernatant was added for the anti-Fc ELISA and 20 µl of supernatant for the anti-His ELISA. After 1 hour incubation at 4°C, CoMiX-Fc were detected with a goat anti-human IgG Fc HRP pAb (Abcam, #ab97225), whereas CoMiX-FHR4 molecules and V_H_H controls were detected with a mouse anti-His HRP-conjugated mAb (Sigma-Aldrich). The plate was revealed with OPD/H_2_O_2_ HRP-chromogenic mix substrate and the absorbance was measured at 492 nm. The clones with the highest production were selected and expanded in cellSTACKs® (Corning) in complete DMEM medium with 10% FBS, Penicillin/Streptomycin and L-Glutamine.

### Flow cytometry

The ability of CoMiX to activate the complement cascade through C3b and C5b9 deposition was evaluated by flow cytometry. HER2-positive BT474 cancer cells were incubated for 1 hour at 4°C with 3-fold serial dilutions of CoMiX molecules, V_H_H controls and therapeutic antibodies, starting from 15 µg/well. In case of molecule combinations, 7.5 µg/well of each was used. Following washing steps, tumor cells were incubated with 25% normal human serum (NHS) diluted in gelatin veronal buffer containing CaCl and MgCl (GVB^++^: 141 mM NaCl, 0.3 mM CaCl_2_, 1 mM MgCl_2_, 0.1% gelatin, 1.8 mM Na-barbital & 3.1 mM barbituric acid, pH 7.3-7.4) for 30 minutes at 37°C. After washing, antibodies were added: mouse anti-human C3/C3b/iC3b mAb (Cedarlane, Sanbio B.V., Uden, The Netherlands) followed by goat anti-mouse IgG AF647 (Invitrogen), mouse anti-human C5b9 (Novus, USA, Centennial) to measure MAC formation, and Live/Dead (L/D) staining (Invitrogen) to quantify complement-dependent cytotoxicity (CDC). The samples were fixed with 1% PFA and analysed with a BD LSR Fortessa™ cell analyzer. The percentage of dead cells was calculated by dividing the number of L/D-positive cells by the total number of cells analyzed. To determine whether CoMiX molecules exert their complement-activating properties through the classical or alternative pathway, both GVB^++^ and GVB^+^ (gelatin veronal buffer containing 3 mM MgCl_2_ and 5 mM EGTA without Ca^2+^ ions) were used to impair the activation of the classical pathway. To analyze the NK-activating properties of CoMiX, we incubated BT474 cells with 5-fold diluted CoMiX molecules starting from 20 µg/condition for 1 hour at 4°C. After washing, cells were co-incubated with human CD16-expressing NK cells (NK92^humCD^^16^) for 4 hours at 37°C and the 421™ anti-human CD107a (Biolegend) antibody as a degranulation marker. GolgiStop™ (BD Biosciences) and GolgiPlug™ (BD Biosciences) were added after 1 hour of incubation.

### CoMiX-mediated cellular phagocytosis of BT474 cells

M2 Macrophages were derived from monocytes of Peripheral blood mononuclear cells (PBMCs) from healthy donors (Luxembourg Red Cross) as previously described ^13, 14^. M2 macrophages were plated into Lab-Tek 8-chamber slides (Nunc, Thermo Fisher Scientific) and were stained with PKH26 (Sigma-Aldrich) forty eight hours later. BT474 cells were stained with CFSE (Invitrogen-Thermo Fisher) and then incubated for 30 min with 20 µg/ml of controls, CoMiX, trastuzumab, pertuzumab or their combination in GVB++ supplemented with 25% C5-deficient NHS to prevent lysis. CFSE-stained complement-coated tumour cells were then washed and added to the KHP26-stained M2 macrophages (ratio was 2:1) and incubated for 18 h at 37°C/5% CO2 in complete RPMI medium. Cells were fixed with 1% PFA, stained with DAPI and analysed with confocal microscopy on the CSU-W1 Spinning Disk/High Speed Widefield confocal microscope lens X4. Three series of 10 pictures were taken for each experimental condition. Phagocytosis was evaluated on 400 macrophages for each condition.

### In vivo models of human BT474 breast cancer xenografts in NUDE mice

The protective capacity of CoMiX molecules was determined *in vivo* with xenografts of human HER2 BT474 expressing cells in NUDE mice. All animal experiments were approved by the animal welfare committee of the Luxembourg Institute of Health and the Ministry of Agriculture of Luxembourg (protocol DII-2018-15). Five weeks old female BALB/c NUDE mice with an average body weight of 25 g were anesthetized with isoflurane and injected subcutaneously with 5×10^6^ Matrigel-mixed (1:1) trastuzumab-sensitive BT474 cells into the mammary fat pads. Tumor growth was measured with calipers and tumor volumes were calculated using the following formula: Volume = (Length x Width x Width)/2. After tumors reached 60 mm^3^ (in 30-40 days), the mice were randomly divided into 12 groups of 5 mice/group to analyze the effect of each CoMiX, the V_H_H(T) control, trastuzumab, pertuzumab and their combinations *in vivo*. The mice were treated 10 times for 10 consecutive days with 100 µg of each CoMiX-FHR4, each CoMiX-Fc, the V_H_H(T) control, trastuzumab, and pertuzumab. Molecule combinations were also tested by injecting 50 µg of each CoMiX-FHR4, CoMix-Fc and each therapeutic antibody. Tumors were measured with calipers every second or third day for 25 days (or until humane endpoints were met), and tumor volumes were recorded. The effects of the most efficient CoMiX [FHR4/V_H_H(T), FHR4/V_H_H(P)] were further explored *in vivo* with trastuzumab-resistant BT474 cells (ATCC CRL-3247) as previously performed with trastuzumab-sensitive BT474 cells. Tumor volume was measured for 37 days or until reaching the humane endpoint as previously described.

### Immunohistochemistry and magnetic resonance imaging

Tumors of sacrificed mice were also collected for immunohistochemistry. To analyse complement activation on xenografts, a single dose of 100 µg of the combination of CoMiX-FHR4 [50 µg FHR4/V_H_H(T) + 50 µg FHR4/V_H_H(P)], of the combination of CoMiX-Fc [50 µg V_H_H(T)/Fc + 50 µg V_H_H(P)/Fc], of 100 µg of the V_H_H(T) control or the combination of 50 µg of each commercial antibody was injected intravenously into the lateral tail vein. One or 6 hours after injection, mice were sacrificed by cervical dislocation. The tumors were surgically removed, placed in OCT cryomold and instantly frozen in liquid nitrogen. Tumors were sectioned to a thickness of 5 µm using the rotary microtome cryostat machine (Leica). A circle drawn with a hydrophobic pen around the tissue section was used to provide surface tension. Tumor sections were fixed with 75% acetone + 25% water cooled at −20°C and dried at room temperature. The sections were blocked with 5% BSA-PBS solution at room temperature for 1 hour and washed 3 times with 1% BSA-PBS between each incubation step. After blocking, CoMiX-FHR4 and V_H_H control sections were incubated with 1 µg of rabbit anti-His mAb (Bethyl) or rabbit anti-human C3d polyclonal Ab (Agilent) primary antibodies, followed by the goat Anti-Rabbit IgG Fc AF568 (Abcam) secondary antibody. The polyclonal rabbit anti-human C3d cross-reacts with mouse complement and is therefore suitable to detect complement deposition in mouse tumors. CoMiX-Fc molecules were revealed with goat anti-human IgG AF647-conjugated antibody (Abcam), which cross-reacts with mouse IgGs. Finally, the sections were fixed with 1% PFA and stained with 200-fold diluted DAPI (Invitrogen). The slides were read on the CSU-W1 Spinning Disk/High Speed Widefield confocal microscope lens X40. Twenty five (5×5) planes were pictured for each tumor section and the results were reconstructed using the NIS Elements Nikon software. To evaluate leukocyte infiltration, BT474 xenografts were collected just after the end of treatment at D+11 in three mice treated with the combination of CoMiX-FHR4 [FHR4/V_H_H(T) + FHR4/V_H_H(P)], trastuzumab or PBS to make cryosections. Cryosections were stained with a monoclonal rat IgG2a anti-mouse NKp46/NCR1 (R&D Systems, clone # 29A1.4) and revealed using a donkey AF568-conjugated anti-rat pAb (Abcam). The sections were fixed and analyzed as described above. Magnetic Resonance Imaging (MRI) was performed on several representative mice of each group. The mice were sacrificed at Day 11 after treatment, and the entire mice were placed in tubes containing cold solution of 4% paraformaldehyde (PFA). The tubes were then left at 4°C until the MRI scan. MRI was performed on a 3T preclinical horizontal bore scanner (MR Solutions, Guilford, UK), equipped with a quadrature volume coil designed for rodent head imaging, with a 17 cm horizontal bore (figure 5C). On the scan day, the heads were placed in a tube with fluorinert (3M, MN, USA). Anatomical FSE T2w (Spatial Resolution: 0.1511×110.1511×111mm; Number of Slices: 30; TE\TR: 68\700011ms; Number of Averages: 1) were used to calculate the tumor volumes per animal. The tumor volume was calculated by the ImageJ software^15^.

### Statistical analyses

Statistical analysis was performed using GraphPad Prism version 10.0.0 for Windows (GraphPad Software, Boston, Massachusetts USA). For all *in vitro* experiments, multiple groups receiving the different single CoMiX molecules or combination treatments were compared using one-way ANOVA and post-hoc Tukey’s tests. For studies in mice, an appropriate sample size (n = 5) was calculated during the study design to obtain groups with a difference of tumor growth of 20% by taking into account a common standard deviation of 5% using a bilateral Student’s T test based on a 95% confidence level. Mice with different tumor sizes were randomized between the groups. The groups were compared using unpaired Student’s T test. A p value < 0.05 was considered as statistically significant (*p < 0.05, **p < 0.01, ***p < 0.001, ****p < 0.0001).

## Results

### Molecular pattern of the different CoMiX

We used the oligomerization scaffold of the C-terminal domain of the α-chain of the C4b-binding protein (C4bp) to enable the formation of hexa- and heptamers. As described in figure 1, Fc-based complement-activating multimeric immunotherapeutic complexes (CoMiX-Fc) display a multivalent targeting function derived from trastuzumab (V_H_H(T)) or pertuzumab (V_H_H(P)), a monovalent tracking function (eGFP) and three dimeric C4bp alpha Fc regions engineered with a dual hinge region between the C4bpα-scaffold and the IgG1 CH2-CH3, ensuring the complement activating properties of the molecule (effector function): V_H_H(T)/Fc and V_H_H(P)/Fc. To evaluate the importance of the Fc dimers, Fc-based multimers without hinge (V_H_H(T)/Fc Δhinge) were also generated as a control for V_H_H(T)/Fc (figure 1). FHR4-based CoMiX molecules (CoMiX-FHR4) include the FHR4 effector function instead of the Fc regions without hinge and the targeting function V_H_H(T) or V_H_H(P). The V_H_H(T) molecule without effector functions was also produced as a control for all CoMiX.

**Figure 1.**
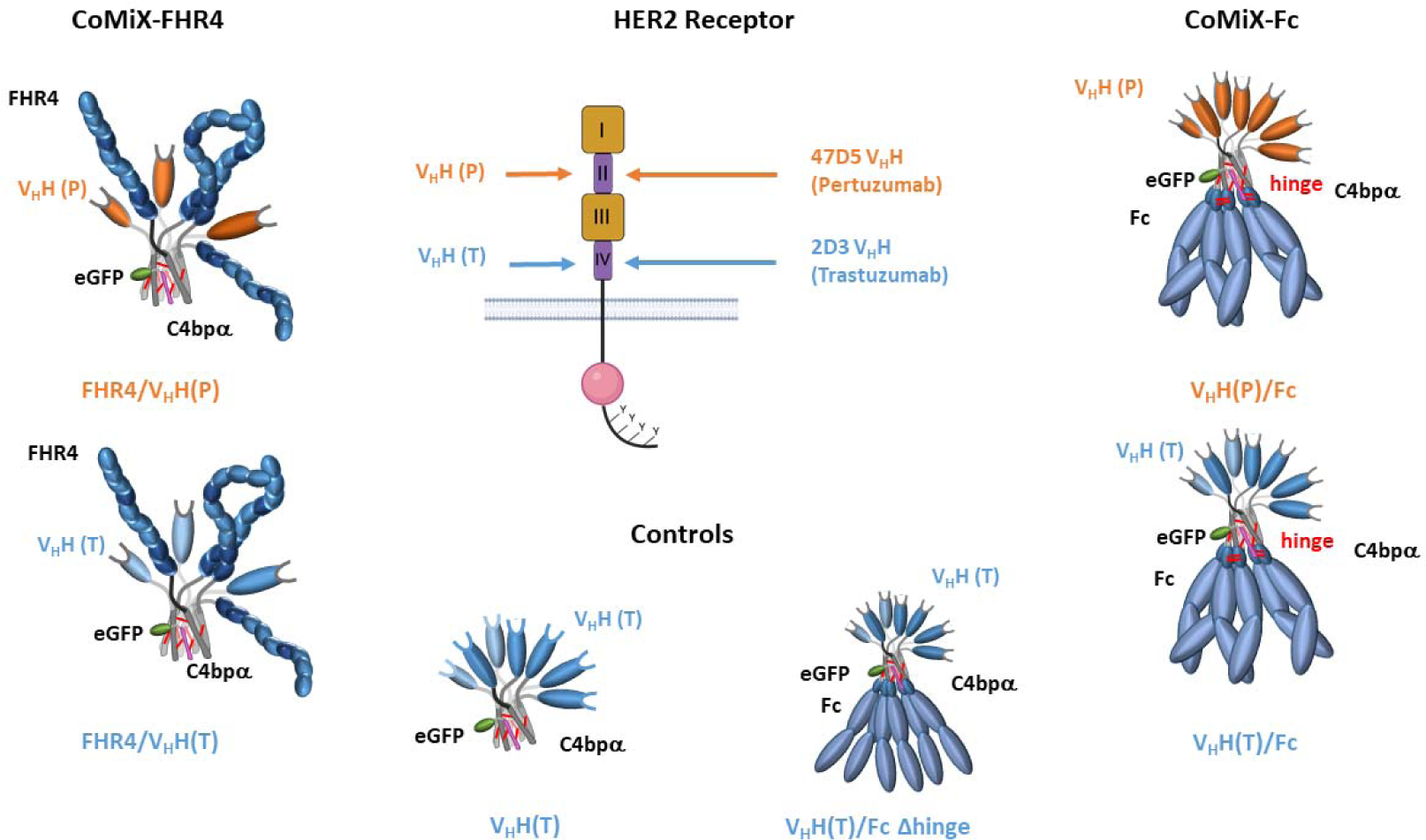
: Design of CoMiX-FHR4, CoMiX-Fc and the controls V_H_H(T) and V_H_H(T)/Fc Δhinge. All constructs are co-transfected with the eGFP.C4bpβ construct, leading to the covalent association of a single eGFP tracking function with the multimeric fusion C4bp α core with the a-chains represented in red lines. We used V_H_H(T) and V_H_H(P), recognising trastuzumab- or pertuzumab-competing HER2 epitopes, respectively, to generate 2 types of CoMiX-FHR4 molecules: FHR4/V_H_H(T) and FHR4/V_H_H(P)] or 2 types of CoMiX-Fc molecules :V_H_H(T)/Fc or V_H_H(P)/Fc. A dual hinge region between the C4bpα-scaffold and the IgG1 CH2-CH3 (represented by two red bands between the two Fc fragments) allows the formation of interchain disulfide bonds and the dimerisation of Fc-regions. C4bpα.His8x V_H_H(T) is the control multimeric molecule with no effector function, and so called V_H_H(T), and V_H_H(T)/Fc Δhinge is the control molecule of V_H_H(T)/Fc without hinge that allows the formation of triple Fc dimers.

The molecular pattern of the produced multimeric immunoconjugates was analyzed under non-reducing (supplementary figure 1AB) and reducing conditions (supplementary figure 1C) by Western blot. Under non-reducing conditions, the FHR4/V_H_H(T) and FHR4/V_H_H(P) molecules display seven bands representing the number of FHR4 molecules in the complexes (supplementary figure 1A and 1B). For CoMiX Fc, the goat anti-human IgG antibody cross-reacts with the V_H_H region of the multimers (supplementary figure 1B). Supplementary figure 1C shows the different monomers under reducing conditions. All molecules have a tracking function (eGFP.C4bpβ) with a size of 50 kDa and a V_H_H anti-HER2 targeting function derived from trastuzumab (V_H_H(T)) or pertuzumab (V_H_H(P)), with a size of 40 or 30 kDa, respectively. The targeting function in CoMiX-Fc molecules is fused together with the effector Fc-function (V_H_H(T).C4bpα.Fc). CoMiX-FHR4 molecules are composed of 120 kDa FHR4.C4bpα.His8x monomers.

### Multimeric immunotherapeutic complexes promote C3b deposition, MAC formation and direct killing of BT474 tumor cells

The effect of the multimeric immunotherapeutic complexes on complement activation was first analyzed by measuring C3b and C5b9 depositions as well as CDC on the HER2-expressing BT474 tumor cell line using flow cytometry (Supplementary figure 2). The terminal complex protein C5b9 was used to measure MAC formation at the surface of BT474 cells. CoMiX-Fc and CoMiX-FHR4 molecules were tested individually or in combination using 3-fold serial dilutions ranging from 15 µg to 0.5 µg/well in case of individual molecules, and from 7.5 µg to 0.25 µg/well of each in case of molecule combinations, in the presence of 25% of NHS and showed a dose-response stimulation of each complement marker and CDC (Supplementary figure 3). The combination of CoMiX molecules enhanced significantly C3b, C5b9 depositions and CDC as compared to the single administrations.

At their highest concentration (figure 2), CoMiX-FHR4 (FHR4/V_H_H(T) and FHR4/V_H_H(P)) led to the highest C3b deposition and CDC, both individually and in combinations (p< 0.0001). CoMiX-Fc (V_H_H (T)/Fc or V_H_H(P)/Fc) were less efficient to facilitate C3b deposition and CDC, however their effect still surpassed the very low complement activating capacity of trastuzumab and pertuzumab Abs (figures 2A and 2C). C5b9 deposition-mediated by CoMiX-FHR4 and CoMiX-Fc harboring V_H_H(T) was more potent than their counterpart harboring V_H_H(P) (figure 2B) (p< 0.0001); nevertheless all molecules exceeded the C5b9 deposition induced by trastuzumab and pertuzumab (p< 0.05). Combinations of CoMiX-FHR4 with CoMiX-Fc enhanced significantly C3b and C5b9 depositions and CDC as compared to CoMiX-Fc alone (p< 0.0001). Overall, CoMiX with the V_H_H(T) targeting system activate the complement system more efficiently than their V_H_H(P) counterparts. The two commercial antibodies have no complement activating effect individually. However, they show increased C3b, C5b9 deposition and CDC when combined. The control V_H_H(T) induces a basal CDC of 20% to the same extent than CoMiX V_H_H(T)/Fc or V_H_H(P)/Fc. To better visualize the effect of each CoMiX and combinations, the correlation between C3b deposition and the percentage of dead cells at 15 µg/well of molecules was depicted on figure 2D. Trastuzumab and pertuzumab have little to no effect on C3b deposition and cell death when used individually. However, they show increased C3b deposition and slight increase in cell death when combined. CoMiX-FHR4 molecules (FHR4/V_H_H(T) and FHR4/V_H_H(P)) demonstrated the highest complement activating properties. CoMiX-Fc molecules (V_H_H(T)/Fc and V_H_H(P)/Fc) have a similar effect on complement activation as the combination of the two therapeutic mAbs. The killing activities of the different combinations with CoMiX-FHR4 and CoMiX-Fc exceeded the individual capacity of all molecules, except for FHR4/V_H_H(T). Taken together, these data indicate that FHR4/V_H_H(T) are the most efficient CoMiX in facilitating C3b deposition and direct killing of tumor cells.

**Figure 2:**
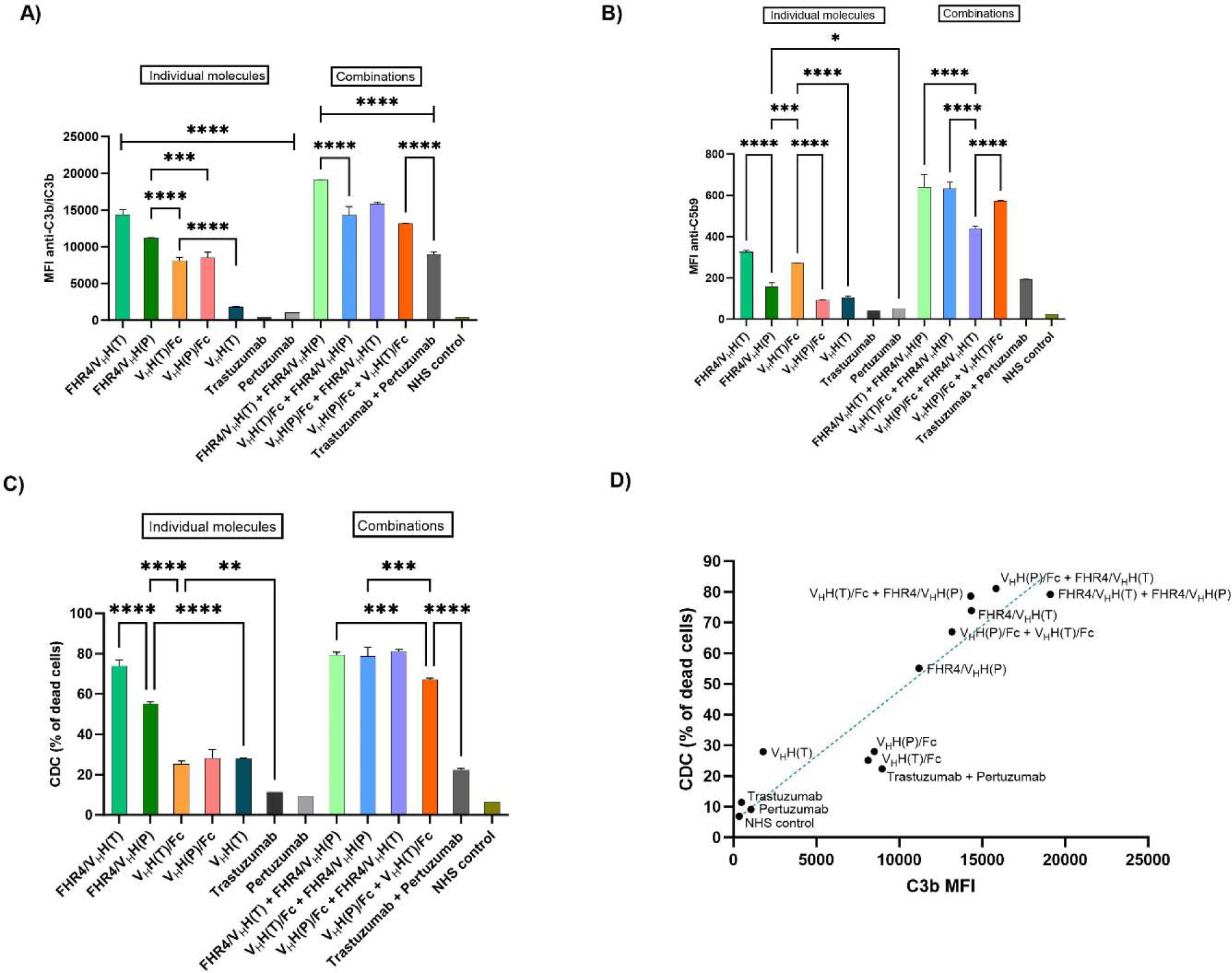
Flow cytometry analysis of C3b/iC3b deposition **(A),** membrane attack complex formation **(B)**, and complement-dependent cytotoxicity **(C)** on BT474 tumor cells incubated with 15 µg/well of multimeric immunotherapeutic complexes. **(A)** C3b/iC3b deposition was detected with mouse anti-human C3b mAb and a secondary goat anti-mouse IgG Ab conjugated with AF647. **(B)** MAC formation was analyzed using anti-C5b9 mAb followed by PE-conjugated anti-mouse IgG pAb. MAC-formation was highest when CoMiX-Fc and CoMiX-FHR4 molecules were combined. **(C)** The percentage of dead cells was calculated by dividing the number of live/dead-positive (dead) cells with the total number of analyzed cells. **(D)** A linear correlation between C3b deposition (MFI) and the percentage of dead cells at 15 µg/well of molecules was observed. Consistent with C3b deposition and MAC-formation, CoMiX-Fc and CoMiX-FHR4 significantly increased the percentage of dead cells compared to control multimers and therapeutic antibodies. Data are presented as mean values ±SD of n = 3 independent experiments. Statistical analysis was performed using a one-way ANOVA and post-hoc Tukey test (**p < 0.005, ***p < 0.001, ****p < 0.0001).

### Mechanisms of action of CoMiX-FHR4 and CoMiX-Fc

In order to assess whether the different types of CoMiX molecules exert their complement activating effect via the classical or alternative pathway, we used 25% NHS diluted in either GVB^++^ or GVB^+^ buffer. GVB^++^ buffer contains Ca^2+^ and Mg^2+^ ions, therefore all three pathways of the complement cascade can be initiated. On the other hand, the absence of Ca^2+^ ions from the GVB^+^ buffer selectively inhibits the classical and lectin pathways, hence only the alternative pathway is active. C3b deposition and the percentage of dead BT474 cells (CDC) were measured by flow cytometry (figure 3). As expected, C3b deposition and the percentage of dead cells are significantly reduced in GVB^+^ buffer compared to GVB^++^ in all cases (p< 0.0001). Incubating BT474 cells with 15 µl/well of FHR4-based and Fc-based CoMiX molecules followed by the addition of normal human serum (NHS) diluted in GVB++ buffer resulted in a 6.5-fold increase in C3b deposition (figure 3A) and a 20 to 30-fold increase in the percentage of dead cells (figure 3B) compared to trastuzumab and pertuzumab. Interestingly, the complement activating properties of FHR4-based CoMiX molecules are reduced only by half in GVB^+^ buffer, whereas the other Fc-based CoMiX molecules and reference mAbs completely lose their activating effect (figure 3CD). Thus, FHR4-based CoMiX molecules facilitate alternative pathway activation, while Fc-based molecules and therapeutic antibodies act as classical pathway activators.

**Figure 3.**
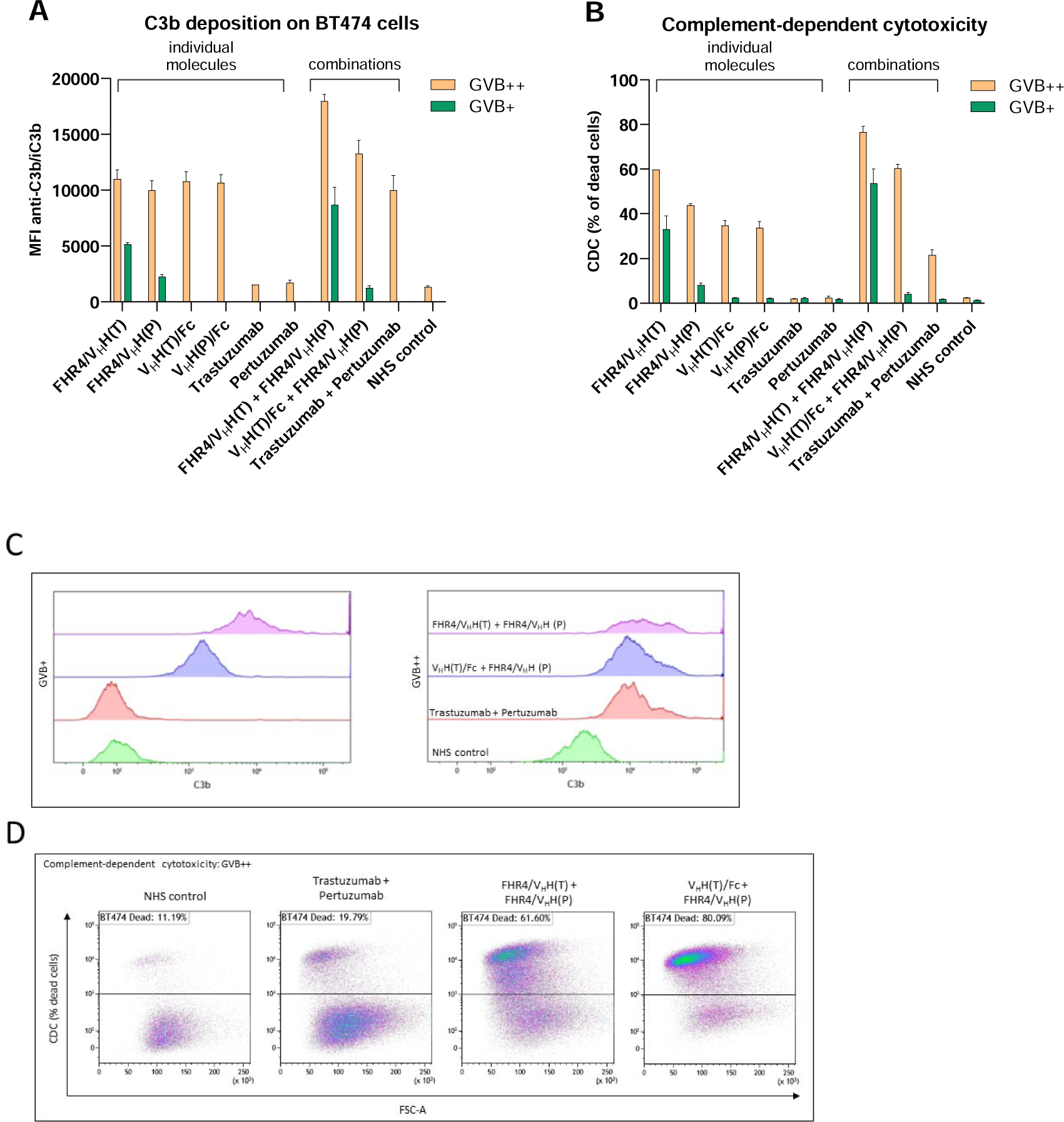
: FHR4-based CoMiX molecules activate the alternative complement pathway, whereas Fc-based CoMiX molecules facilitate classical pathway activation. C3b deposition **(A)** and CDC **(B)** of BT474 tumor cells incubated with saturating concentrations (15 µl/well) of CoMiX molecules and control mAbs individually or in combinations. 25% NHS diluted in either GVB^++^ or GVB^+^ buffer was added for 30 minutes at 37°C. Inhibition of the classical complement pathway by using GVB^+^ buffer completely disrupts the complement activating properties of Fc-based CoMiX molecules, trastuzumab and pertuzumab. Data are presented as mean values ±SD of n = 3 independent experiments. Statistical analysis was performed using a two-way ANOVA test between GVB^++^ and GVB^+^ conditions for each molecule. All comparisons between GVB^++^ and GVB^+^ reached statistical significance (****p < 0.0001). **(C)** Representative histogram plots on live BT474 cells of C3b MFI for the combinations of molecules with GVB^+^ and GVB^++^ conditions are shown. **(D)** Representative dots plots of live and dead BT474 cells with the different combinations are depicted.

NK cells bind IgG antibodies though their FcγRIIIa receptors (CD16) and facilitate the destruction of opsonized target cells via ADCC. As Fc-based multimeric immunotherapeutic complexes have several Fc regions in their structure, we hypothesized that these molecules could be effective NK cell activators. HER2-positive BT474 tumor cells were incubated with CoMiX and the human CD16-expressing NK cell line (NK92^humCD16^) for 4 hours, and NK cells were analyzed by flow cytometry (figure 4 and supplementary figure 4). The expression of the degranulation marker CD107a was highly upregulated by V_H_H(P)/Fc and trastuzumab and half less by V_H_H(T)/Fc (p <0.0001, figure 4A). CoMiX-FHR4 molecules having no Fc region did not induce significantly CD107a expression similarly to V_H_H(T) or pertuzumab (figure 4A). In concordance, IFN-γ expression was significantly enhanced by V_H_H(T)/Fc and V_H_H(P)/Fc, as well as trastuzumab and pertuzumab (p < 0.0001, figure 4B) as compared to the CoMiX-FHR4 molecules and controls. Without hinge (V_H_HT/Fc Δhinge) the percentage of CD107a-positive NK cells is reduced by 16% when compared to CoMiX-Fc molecules. These data show that the hinge region is mandatory for the assembly of functional CoMiX-Fc molecules with multiple Fc-dimers, capable of efficiently binding to Fcγ receptors on NK cells (figure 4C). The Fc regions of trastuzumab and pertuzumab evoke an overall similar capacity to trigger NK function as their CoMiX-Fc molecules counterparts, although pertuzumab showed a decrease capacity of degranulation as compared to CoMiX V_H_H(P)/Fc (p < 0.0001). In conclusion, CoMiX-FHR4 use the alternative complement pathway whereas the presence of the IgG1 hinge region between the C4bpα-scaffold and the IgG1 CH2-CH3 is crucial to the Fc fragment to activate NK cells in the presence of BT474 target cells.

**Figure 4:**
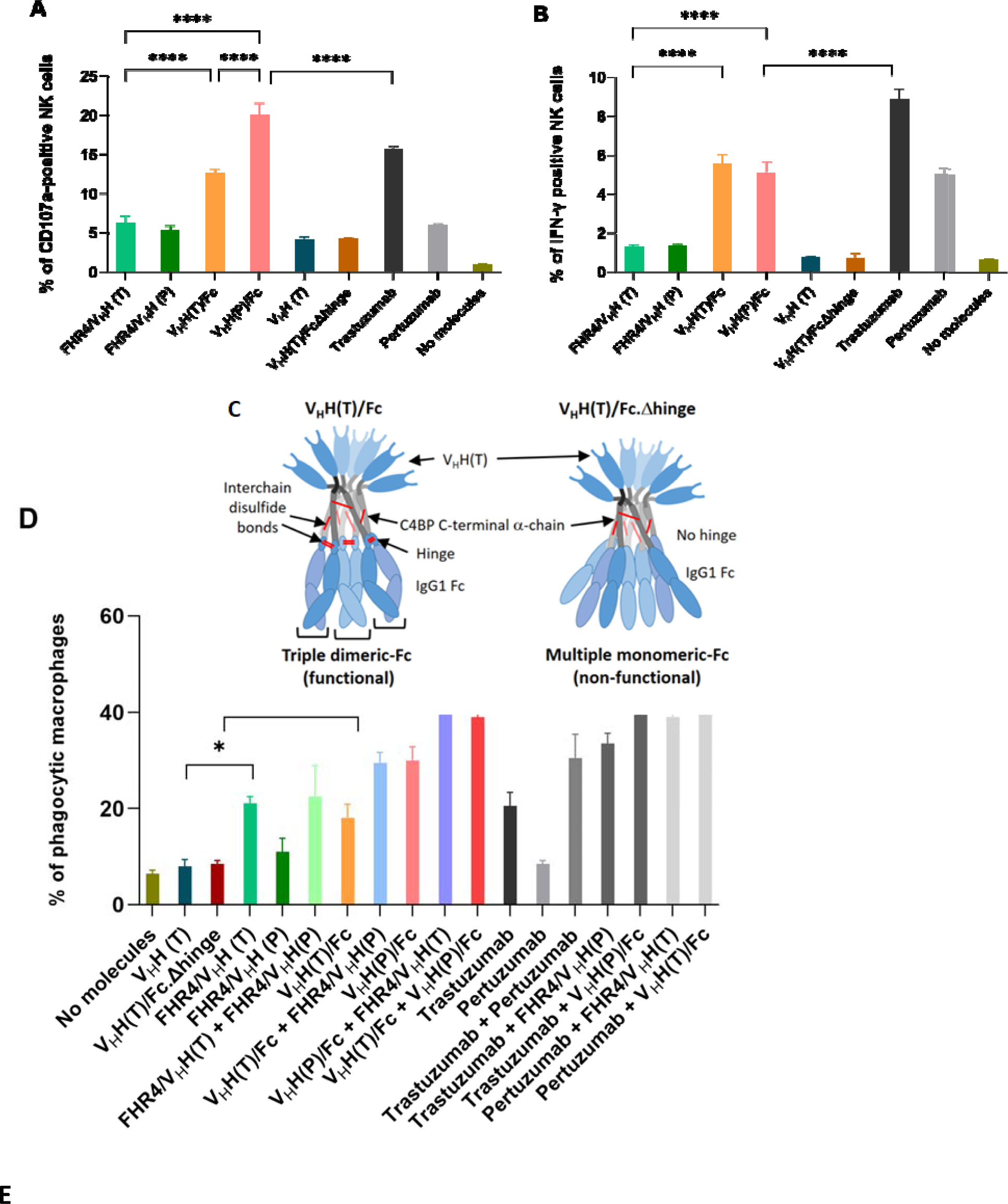
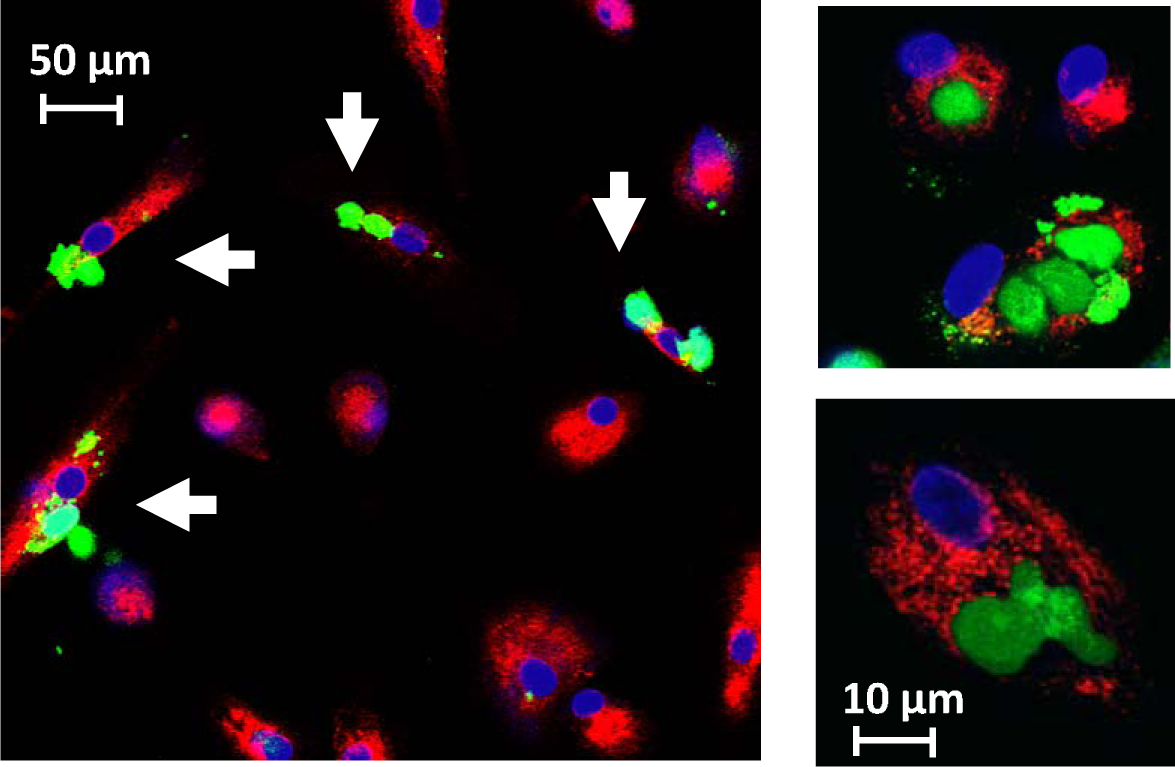
CoMiX-Fc enables NK cell activation and BT474 phagocytosis mediated by M2 macrophages. **(A)** HER2-positive BT474 tumor cells incubated with 15 μg/well of CoMiX molecules were co-incubated with NK92^humCD^^16^ cells. The expression of the CD107a degranulation marker was analyzed by flow cytometry. **(B)** The intracellular accumulation of the cytokine IFN-γ was likewise analyzed by flow cytometry. Data are presented as mean values ±SD of the triplicates of one representative experiment of three independent experiments. Statistical analysis was performed using a one-way ANOVA and post-hoc Tukey test (***p < 0.001). **(C)** Graphic representation of the functional triple dimeric V_H_HT/Fc and its non-functional control V_H_HT/Fc Δhinge. Without hinge, functional Fc-dimers were unable to assemble, forming instead multiple monomeric Fc’s, resulting in a non-functional CoMiX-Fc molecule. **(D)** Complement-dependent macrophage-mediated phagocytosis of human BT474 cells. CFSE-stained BT474 tumor cells were incubated with controls, CoMiX, trastuzumab + pertuzumab or their combination with 25% C5-deficient human serum to prevent lysis. Percentage of phagocytic macrophages was measured. Data are presented as mean values ± SD out of n = 3 series of 10 confocal images for each condition. Statistical analysis was performed using a one-way ANOVA and post-hoc Tukey test (*p < 0.05, **p < 0.01). **(E)** Left: Confocal microscopy images of BT474 phagocytosis mediated by M2 macrophages when incubated with CoMiX V_H_H(T)/Fc. The white arrows show the phagocytic macrophages (red) having engulfed BT474 cell(s) and harboring a green or yellowish color. Scale bar represents 50 µm. Right: Details of a phagocytic macrophage (red) at higher magnification (60× oil immersion). Scale bar represents 10 µm.

We previously showed that FHR4 induced complement-dependent macrophage-mediated phagocytosis on BT474 tumor cells^13^. We therefore compared the effects of CoMix-Fc to CoMiX-FHR4, trastuzumab, pertuzumab and their combinations on macrophage-mediated phagocytosis using confocal microscopy and C5-deficient serum to inhibit complement-dependent killing (figure 4DE). As previously described, CoMiX-FHR4 V_H_H(T) enhanced phagocytosis of BT474 target cells to the extent of trastuzumab (around 20%, p <0.05). CoMiX V_H_H(P)/Fc increased the number of BT474 cells phagocytosed by macrophages up to 30% (p < 0.005) but not CoMiX V_H_H(T)/Fc. All combinations between CoMiX-Fc, CoMix-FHR4, trastuzumab and pertuzumab enhanced phagocytosis to a mean of around 40% (p < 0.01), with the exception of the trastuzumab/pertuzumab and trastuzumab/FHR4 V_H_H(P) combinations. We here confirmed that the Fc/FcγRs interactions are functional in CoMiX-Fc to stimulate phagocytosis mediated by macrophages and that the hinge region is mandatory for the assembly of functional CoMiX-Fc molecules with multiple Fc-dimers.

### CoMiX-FHR4 and CoMix-Fc delay tumor progression in vivo by activating the complement

Tumor xenograft animal models are widely used in preclinical studies to evaluate the efficacy of novel anticancer therapies. The effect of CoMiX molecules, the controls and the commercial therapeutic antibodies trastuzumab and pertuzumab was analyzed in vivo using the BT474 cell line in a xenograft mouse model. Female BALB/c NUDE mice were injected with trastuzumab-sensitive BT474 cells into the mammary fat pads. As described in figure 5A, when the tumor volumes reached ∼60 mm^3^, the mice were treated for 10 days with 100 µg of CoMiX-FHR4 molecules, CoMiX-Fc molecules, V_H_H(T) controls or monoclonal therapeutic antibodies. Four molecule combinations were also tested, by injecting 50 µg of each. Tumor volumes were measured for 25 days (or until humane endpoints were met) (figure 5B). MRI was performed for several representative mice after sacrifice^15^ and images are depicted in figure 5C. CoMiX-FHR4 molecules reduced the tumor volume by a factor of 7.33 compared to the PBS control. V_H_H(T)/Fc had no effect on tumor growth, while V_H_H(P)/Fc led to a 2.75-fold tumor volume reduction that was superior to the reduction produced by pertuzumab. Trastuzumab and its combination with pertuzumab remained the most potent regimen that completely inhibited tumor growth in the xenograft model.

To confirm the ability of CoMiX-FHR4 molecules to infiltrate the tumor tissue and activate the complement, immunohistochemistry of tumor sections was performed (figure 6). BT474 cells were injected into the mammary fat pads of female BALB/c NUDE mice. After the tumor volume reached ∼60 mm^3^, the mice were injected with 100 µg of CoMiX-FHR4 (50 µg of FHR4/V H(T) + 50 µg of FHR4/V_H_H(P)), or 100 µg of CoMiX-Fc (50 µg of V_H_H(T)/Fc + 50 µg V_H_H(P)/Fc), or PBS, or the 100 µg of the V_H_H (T) control, or the commercial antibodies (50 µg trastuzumab + 50 µg pertuzumab) intravenously. One hour after injection the molecules were detected mainly at the tumor periphery and colocalize with C3b-deposition (left panels). 6 hours after injection, the whole tumor is infiltrated with CoMiX-FHR4 or CoMiX-Fc molecules (upper panels) and complement activation is not restricted to the periphery but occurs inside the whole tumor tissue (middle panels). V_H_H(T) shows a significantly reduced complement activating capacity as compared to CoMiX-FHR4 molecules or CoMiX-Fc molecules 6 hours after injection (lower panel). CoMiX-Fc recruit C1q and activate the classical complement pathway like trastuzumab in combination with pertuzumab.

**Figure 5.**
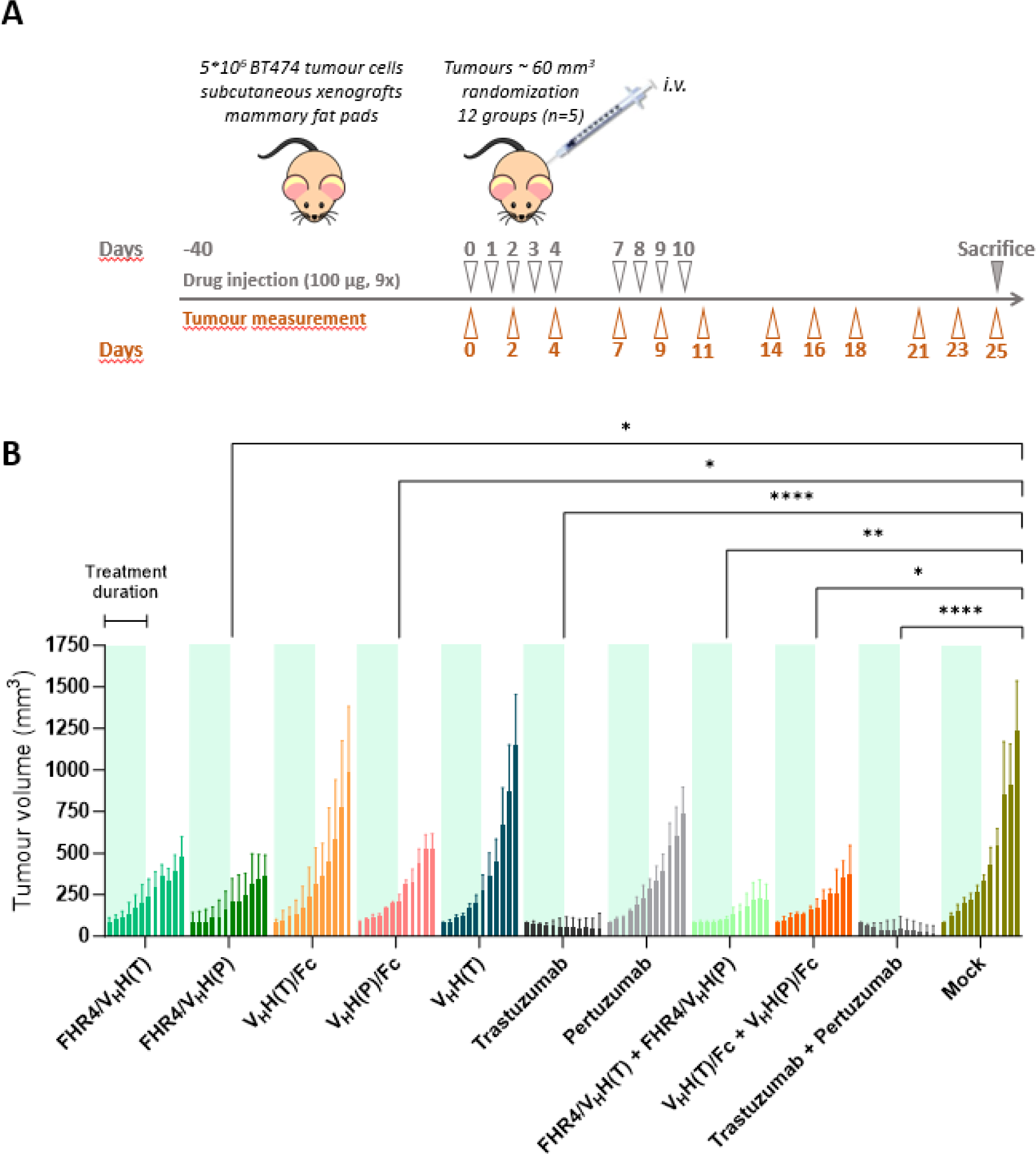
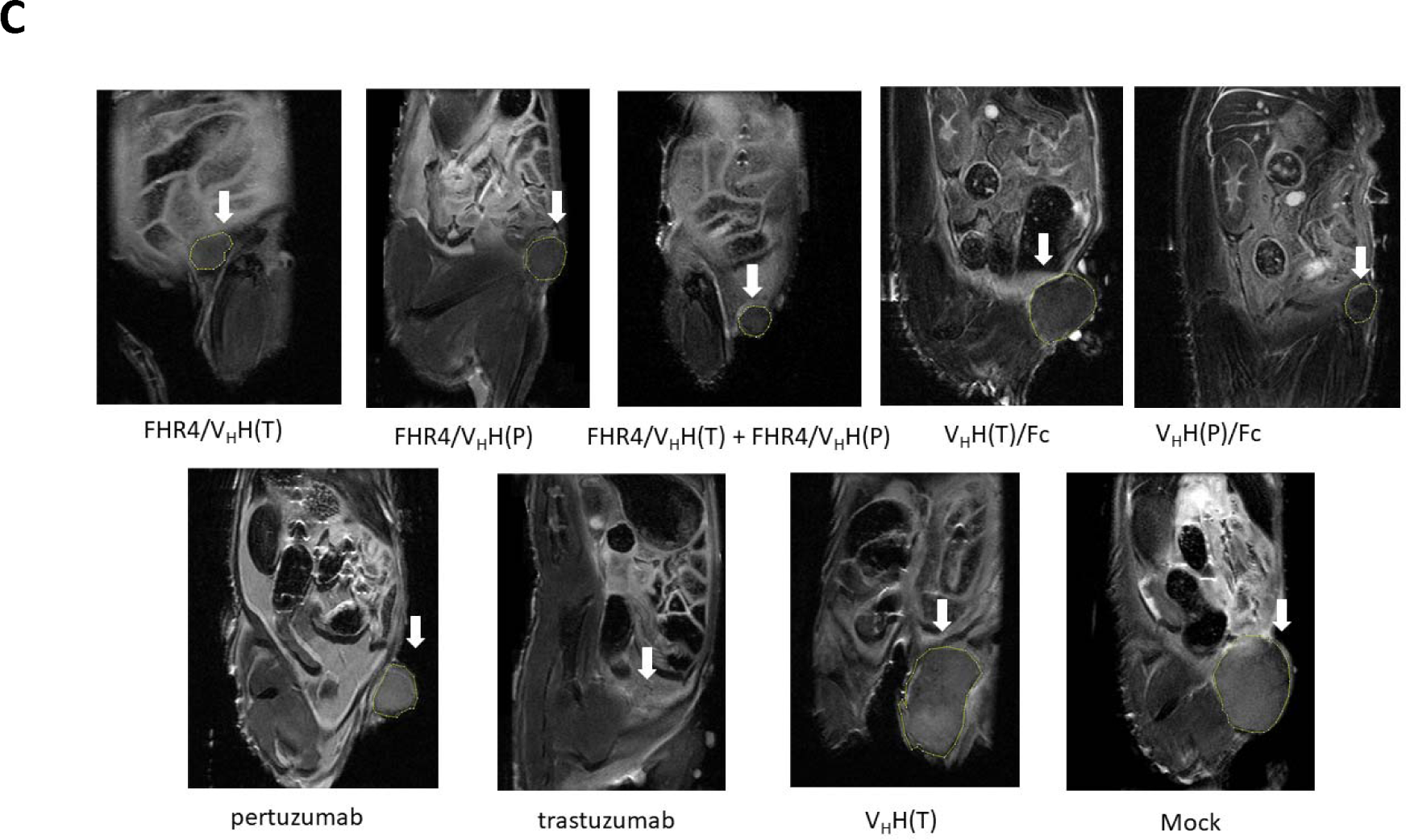
FHR4/V_H_H(T), FHR4/V_H_H(P) and V_H_H(P)/Fc molecules reduce tumor growth, whereas V_H_H(T)/Fc has no beneficial effect. **(A)** Experimental design for the measurement of the subcutaneous xenografts mammary fat pads volume in the presence of different CoMiX molecules and control antibodies. BT474 cells were injected into the mammary fat pads of mice. When the tumor volume reached ∼60 mm^3^, the mice were injected intravenously into the lateral tail vein with 100 µg of CoMiX molecules or control antibodies. The injection was repeated 9 times on days 1, 2, 3, 4, 7, 8, 9 and 10 after the first injection. For molecule combinations, 50 µg of each were injected. The tumors were measured every second or third day with calipers. **(B)** The therapeutic effects of five CoMiX molecules [FHR4/V_H_H(T), FHR4/V_H_H(P), V_H_H(T)/Fc, V_H_H(P)/Fc, V_H_H(T)] and two control antibodies (trastuzumab and pertuzumab) were evaluated individually and in combination. The treatment duration is indicated for all groups in green, as mentioned for the first FHR4/V_H_H(T) group in the left side of the graph. CoMiX-FHR4 molecules are more effective than CoMiX-Fcs, while the molecules that have the pertuzumab-competing epitope are more effective than those with the trastuzumab-competing epitope. **(C)** MRI images of a representative tumor for each group. Mice were sacrificed at the end of the treatment.

**Figure 6:**
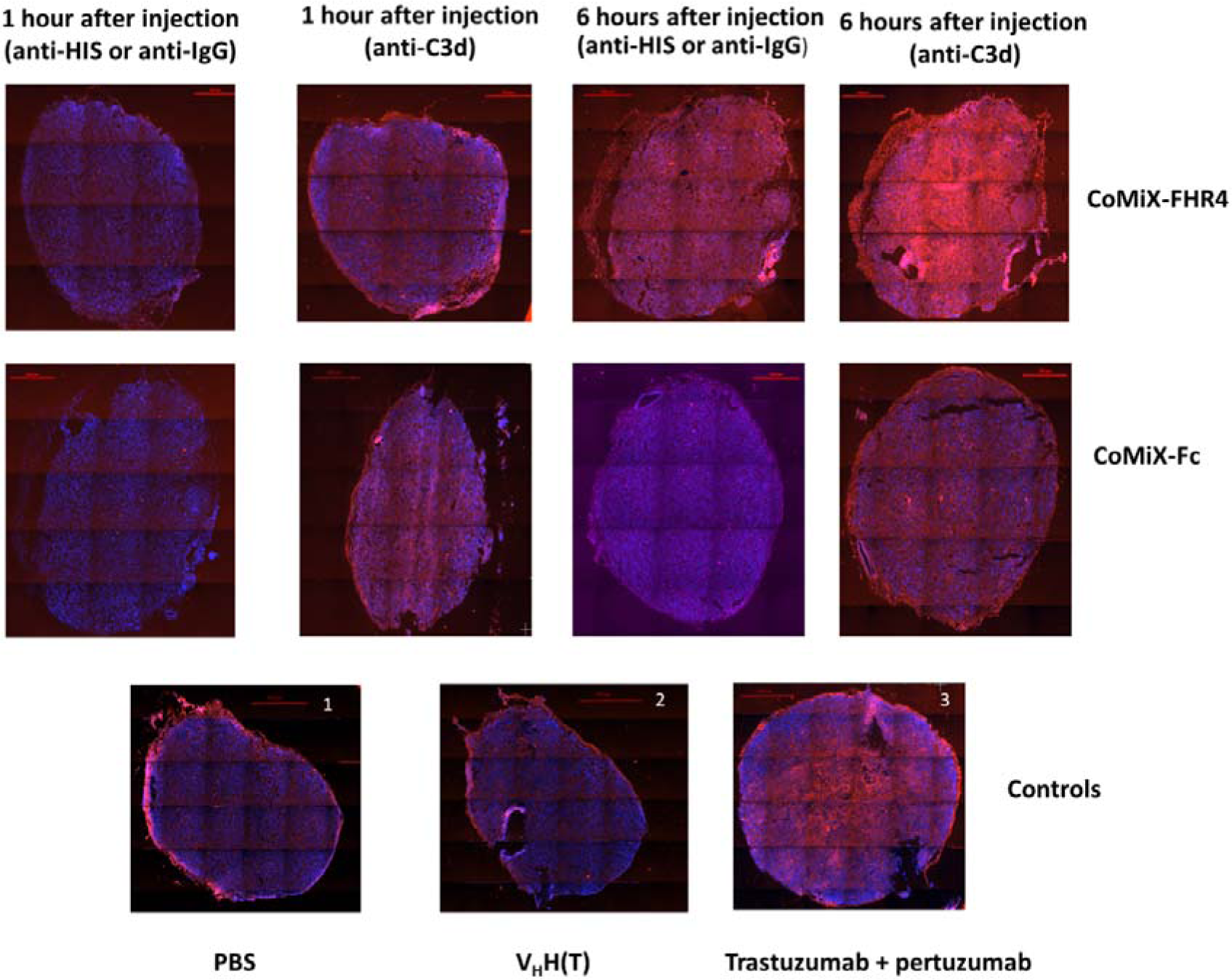
Immunofluorescent staining of tumor sections collected 1 or 6 hours after injection of CoMiX-FHR4 (upper panel), CoMiX-Fc (intermediate panel) or controls (lower panel with anti-C3d staining): PBS (1), V_H_H(T) (2), trastuzumab + pertuzumab (3). CoMiX were visualized with either a rabbit anti-His mAb followed by the goat Anti-Rabbit IgG Fc AF568- or a goat anti-human IgG AF647-conjugated antibody. Complement activation was visualized using the polyclonal rabbit anti-C3d antibody followed by AF568-conjugated anti-rabbit IgG. One hour post-injection, the infiltration of molecules into the tumor tissue is already visible, however complement activation occurs predominantly on the periphery of the tumors. Six hours after treatment, the molecules homogeneously infiltrate the tumor and strong complement activation can be detected throughout the whole tissue. Compared to CoMiX-FHR4 molecules, the V_H_H(T) control (2) shows decreased infiltration and reduced complement activation, present only at the periphery of the tumor, even if collected 6 hours after injection. Trastuzumab + pertuzumab (3) were used as positive controls and showed significant infiltration and complement activation.

Trastuzumab exerts a strong antitumor effect by inhibiting cell growth via the upregulation of the cyclin-dependent kinase inhibitor p27, and the reduction of vascular endothelial growth factor (VEGF) levels^16^. While this is extremely effective at the early stages of treatment, cells quickly develop various resistance mechanisms. The biological activity of CoMiX consists of a rapid activation of the host complement system’s alternate/classic pathways. External recruitment of the complement system is highly unlikely to lead to treatment resistance, in contrast to currently used therapeutic antibodies such as trastuzumab^10^. The effects of CoMiX FHR4/V H(T) and CoMiX FHR4/V_H_H(P) were further explored in vivo (figure 7A) using a trastuzumab-resistant BT474 cell line (ATCC CRL-3247). Injecting trastuzumab for 10 days had no effect on tumor growth, in contrast to the CoMiX-FHR4 molecules which reduced the growth of trastuzumab-resistant cancer cells up to 25 days after treatment initiation. Therefore CoMiX-FHR4 molecules could be an attractive therapeutic strategy when resistance to trastuzumab occurs. BT474 tumors resistant to trastuzumab treated with CoMiX-FHR4, trastuzumab or PBS where collected just after the 2-week treatment at D+11, and processed for immunohistochemistry with an anti-mouse NKp46 antibody (figure 7B). A massive NK cell infiltration was observed in the tumor treated with CoMiX-FHR4, whereas the infiltration was much weaker in the tumor treated with trastuzumab and absent in the PBS-treated tumor. The cell densities in the tumor treated with CoMiX-FHR4 were lower than in the tumor treated with trastuzumab or in the PBS condition, suggesting that the CoMiX molecules are able to trigger a massive infiltration of NK cells within the tumor in contrast to trastuzumab.

**Figure 7.**
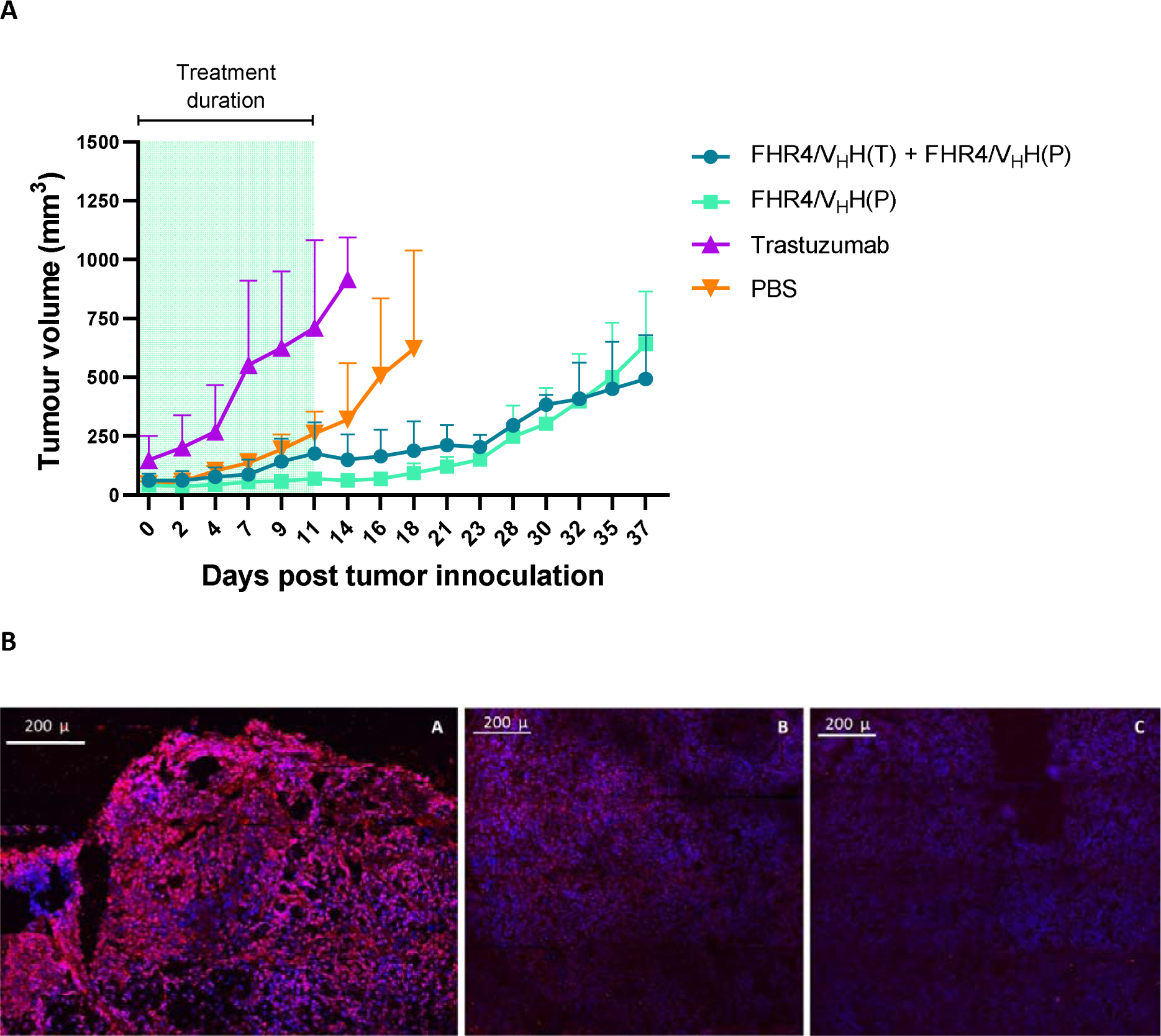
(A) FHR4/V_H_H(T) and FHR4/V_H_H(P) CoMiX molecules exert their anti-tumor effect on trastuzumab-resistant BT474 cells. Trastuzumab-resistant BT474 cells were injected into the mammary fat pads of female BALB/c NUDE mice. When the tumor volume reached ∼60 mm^3^, the mice were injected with 100 µg of CoMiX-FHR4 molecules, trastuzumab or PBS, as described on figure 5A. The tumors were measured every second or third day until day 37 of the study or until meeting a humane endpoint. Trastuzumab and PBS had no beneficial effect on tumor growth, whereas CoMiX-FHR4 molecules were shown to significantly reduce tumor progression. **(B)** Cryosections of trastuzumab-resistant BT474 tumor xenografts collected just after the end of the treatment (at D+11). Tumors were embedded in OCT and snap frozen in OCT. Four micrometer cryosections were made and stained with a monoclonal rat IgG_2A_ anti-mouse NKp46/NCR1 antibody and revealed using a donkey anti-rat AF568-conjugated pAb. Confocal microscope was used to make pictures (lens X40), monitored by the Nikon NIS-Elements software which allowed to assemble pictures to get a large field overview of the tumors. _A)_ tumor treated with combined CoMiX-FHR4 [FHR4/V_H_H(T) + FHR4/V_H_H(P)], _B)_ tumor treated with trastuzumab, _C)_ tumor treated with PBS (mock).

Taken together, our in vivo data are consistent with the data obtained in vitro on BT474 cells. CoMiX-FHR4 and CoMiX-Fc lead to a fast and massive complement deposition and subsequent CDC, but also to the recruitment of innate effector cells such as NK cells. Our approach may offer a new alternative for tumors that became resistant to therapeutic antibodies such as trastuzumab, and, more generally, be an alternative to commonly applied immunotherapeutic approaches of cancer.

## Discussion

HER2-targeted therapies have changed the paradigm for patients and significantly improved their prognosis over the past few decades. For most patients with advanced HER2-positive breast cancer, trastuzumab or pertuzumab are first line treatments. Breast cancer remains nevertheless a clinical challenge due to emerging resistance to HER2 therapies. Therefore, novel therapeutics are still needed. Immunotherapy with directed complement activation is a promising approach^17^. The activation of the classical pathway of complement contributes to the therapeutic efficacy of Rituximab^18–20^. Factor H is a key complement regulator in plasma that can be hijacked by bacteria^21, 22^ or cancer cells to evade complement-mediated cytotoxicity^20, 23^. Factor H related protein 1 to 4 have different activation or inhibition effect. We have previously shown that BT474 tumor cells establish a complement inhibitory threshold, overcome by high valences of FHR4 when directed to the surface of tumor cells^13^. By lacking the regulatory domains of FH, but not the surface binding region of the protein, FHR4 competes with FH for C3b, iC3b and C3d binding^24^. Fc regions, however, mediate several effector functions, including classical complement pathway activation, as well as activation of immune cells (phagocytes, NK cells) via various Fc receptors.

Here, we demonstrated that both CoMiX-FHR4 and CoMiX-Fc molecules have a complement activating capacity on HER2-positive BT474 cancer cells in vitro and in vivo that induce direct lysis of tumor cells. CoMiX-FHR4 molecules [FHR4/V_H_H(T) and FHR4/V_H_H(P)] had a superior effect in vitro on complement activation and CDC when compared to CoMiX-Fc molecules (V_H_H (T)/Fc or V_H_H(P)/Fc) and therapeutic antibodies (trastuzumab and pertuzumab) (figure 2). The two newly generated CoMiX-Fc molecules (V_H_H(T)/Fc or V_H_H(P)/Fc) also increased complement activation on cancer cells compared to their therapeutic antibody counterparts that use the same HER2 epitope, even though less effectively than CoMiX-FHR4. We showed here that, by competing with FH, CoMiX-FHR4 molecules serve as a platform for the assembly of the alternative pathway C3-convertase whereas CoMiX-Fc molecules activate the classical pathway. Therefore, the combination of CoMiX-Fc with CoMiX-FHR4 molecules resulted in enhanced complement activation. Since V_H_H(T) and V_H_H(P) moieties recognise different epitopes, their effect was also additive when two CoMiX-FHR4 or CoMiX-Fc constructs were combined. Furthermore, by measuring the ratio between the percentage of dead cells (CDC) and median fluorescence intensity of C3b deposition, we observed that CoMiX FHR4/V_H_H(T) was the most effective molecule, followed by FHR4/V_H_H(P). Importantly, CoMiX-Fc molecules can activate NK cells and induce, through their Fc regions, complement-mediated phagocytosis of BT474 cells by M2 macrophages (figure 4). V_H_H(P)/Fc was able to activate NK cells and complement-mediated phagocytosis with greater efficacy than pertuzumab, while V_H_H(T)/Fc had a similar effect as trastuzumab.

In a second part of the study, we tested the antitumor growth effect of CoMiX-FHR4 and CoMiX-Fc in a human BT474 cell line-derived xenograft model in NUDE mice. After systemic injection, the diffusion of CoMiX into the tumor was fast, and the tumor demonstrated massive complement activation 6h after injection. FHR4/V_H_H(T), FHR4/V_H_H(P) and V_H_H(P)/Fc molecules reduced tumor growth but not V_H_H(T)/Fc that showed less efficiency for phagocytosis by macrophages and degranulation of NK cells than V_H_H(P)/Fc (figure 4). Trastuzumab was the most potent treatment, indicating that complement activation alone is less efficient than the additive effect of cell signaling inhibition with ADCC/ADCP induced by the therapeutic antibody on limiting tumor growth through the trastuzumab epitope, but not through the pertuzumab epitope. Of note, V_H_H(P)/Fc was superior to pertuzumab for reducing tumor growth in the xenograft model in vivo. This could be explained by its greater efficacy to activate NK cells and complement-mediated phagocytosis. The combined use of two CoMiX-FHR4 recognizing different HER2 epitopes enhanced the C3b deposition, MAC formation, CDC, and therapeutic efficacy in vivo, but was still inferior to trastuzumab’s efficacy alone. However, when tested on a trastuzumab-resistant BT474 cell line, the most potent combination of FHR4/V_H_H(T) and FHR4/V_H_H(P) displayed a strong anti-tumor growth effect.

The complement system has a strong potential to stimulate inflammation. It is, however, tightly regulated, and membrane-anchored complement regulatory proteins, such as CD46, CD55 and CD59, are overexpressed on host cells to prevent high complement activation^25, 26^. Several other soluble complement regulatory proteins are also present in body fluids, such as FH, factor H-like protein 1 (FHL1), classical pathway inhibitors like C4b-binding protein (C4bp), C1 inhibitor as well as terminal pathway inhibitors like clusterin or vitronectin. Therefore, the tight regulation of the soluble complement cascade combined to a directed C3b activation on targeted tumor cells could represent an alternative and safe anticancer therapy^27^. Importantly, complement components are known to regulate the function of the tumor microenvironment by favoring both tumor-promoting and tumor-suppressing responses^1, 11^. Thus, the effects of CoMiX should be further evaluated in this process.

In conclusion, although CoMiX-FHR4 and CoMiX-Fc were not superior to trastuzumab to reduce tumor growth in vivo, as a large percentage of patients easily develop resistance to one or the two conventional monoclonal antibodies, our molecules could delay the emergence of resistance. Since the mechanism of action of CoMiX differs from those of trastuzumab and pertuzumab, and activates the complement system promptly and NK cell infiltration more effectively into the tumor than conventional antibodies, CoMiX FHR4 or CoMiX V_H_H(P)/Fc could be used as second line treatments when resistance to trastuzumab occurs or pertuzumab therapy fails.

## Ethics approval

All animal experiments were approved by the animal welfare committee of the Luxembourg Institute of Health and the Ministry of Agriculture of Luxembourg (protocol DII-2018-15).

## Patient consent for publication

Not required

## Availability of data and statement

All data are available in the main text or the online supplemental materials.

## Author’s Contribution

Conceptualization, X.D., C.S.D., J.H.M.C; methodology, X.D., C.S.D., J.H.M.C, J.M.P, G.I., J.Y.S., G.K., I.B., J.Z.; validation, B.B., G.I., C.S.D., J.H.M.C, J.Z.; analysis, X.D., C.S.D., J.H.M.C, B.B., J.M.P, G.I., J.Y.S., G.K., J.Z.; data curation, J.Z, B.B, C.S.D.; writing—original draft preparation, B.B, C.S.D., J.Z., X.D.; writing—review and editing, B.B, C.S.D., J.M.P, G.I., J.Y.S, J.Z., G.K., I.B., J.Z., J.H.M.C, X.D.; resources, C.S.D, I.B, G.I., J.M.P., J.Y.S, J.Z., X.D.; funding acquisition, C.S.D., I.B. All authors have read and agreed to the published version of the manuscript.

## Acknowledgements

We would like to acknowledge Brigitte Reveil for her technical assistance and Pr Béatrice Clemenceau, University of Nantes for providing the NK92^humCD16^cell line.

## Supplemental Material

**Supplementary figure 1.**
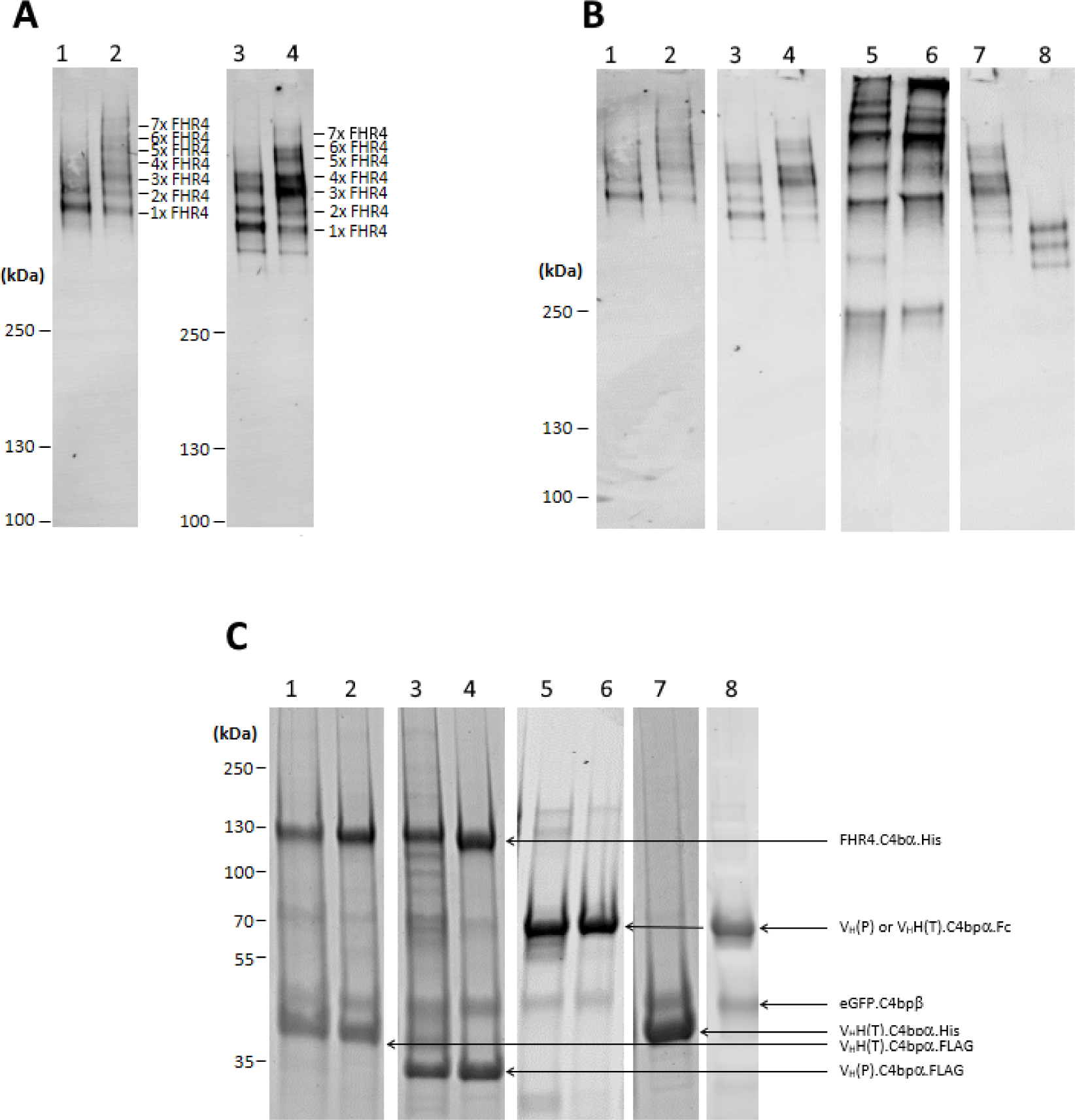
: Visualization of the molecular pattern of purified multimeric immunoconjugates by Western blot analysis of complexes separated under non-reducing conditions. **(A, B)** and SYPRO Ruby protein gel staining under reducing conditions **(C)**. **(A) 1.** FHR4/V_H_H(T) fraction 1, **2.** FHR4/V_H_H(T) fraction 2/3, **3.** FHR4/V_H_H(P) fraction 1, **4.** FHR4/V_H_H(P) fraction 2/3. Under non-reducing conditions, seven bands are visible for the different fractions corresponding to FHR4-valencies varying between 1 and 7. The pooled fractions f2 and f3 display higher FHR4-valencies than their f1 counterpart and were used for further experiments. **(B) 1.** FHR4/V_H_H(T) fraction 1, **2.** FHR4/V_H_H(T) fraction 2/3, **3.** FHR4/V_H_H(P) fraction 1, **4.** FHR4/V_H_H(P) fraction 2/3, **5.** V_H_H(P)/Fc, **6.** V_H_H(T)/Fc, **7.** V_H_H(T)/Fc Δhinge, **8.** V_H_H(T). The different molecular species were analyzed and revealed with a goat anti-human IgG antibody that cross-reacts with the V_H_H region. **(C) 1.** FHR4/V_H_H(T) fraction 1, **2.** FHR4/V_H_H(T) fraction 2/3, **3.** FHR4/V_H_H(P) fraction 1, **4.** FHR4/V_H_H(P) fraction 2/3, **5.** V_H_H(P)/Fc, **6.** V_H_H(T)/Fc, **7.** V_H_H(T)/Fc Δhinge, **8.** V_H_H(T). The multimers were also analyzed by SYPRO Ruby gel staining under reducing conditions. Three bands can be observed for FHR4/V_H_H(T) and FHR4/V_H_H(P), representing the monomeric forms of the FHR4.C4bpα.His (120 kDa), eGFP.SCR3.C4bpβ (50 kDa) and the V_H_H(T).C4bpα.FLAG (40 kDa) or V_H_H(P).C4bpα.FLAG (30 kDa) targeting components. V_H_H(T)/Fc and V_H_H(P)/Fc molecules display two bands representing the eGFP.SCR3.C4bpβ and V_H_H(T).C4bpα.Fc chains and V_H_H(P).C4bpα.Fc, respectively. The V_H_H(T) control molecule has no FHR4- or Fc-effector functions, only targeting (V_H_H(T).C4bpα.His) and tracking (eGFP.SCR3.C4bpβ) functions, whereas V_H_H(T)/Fc Δhinge shows one band for V_H_H(T).C4bpα.Fc.

**Supplementary figure 2.**
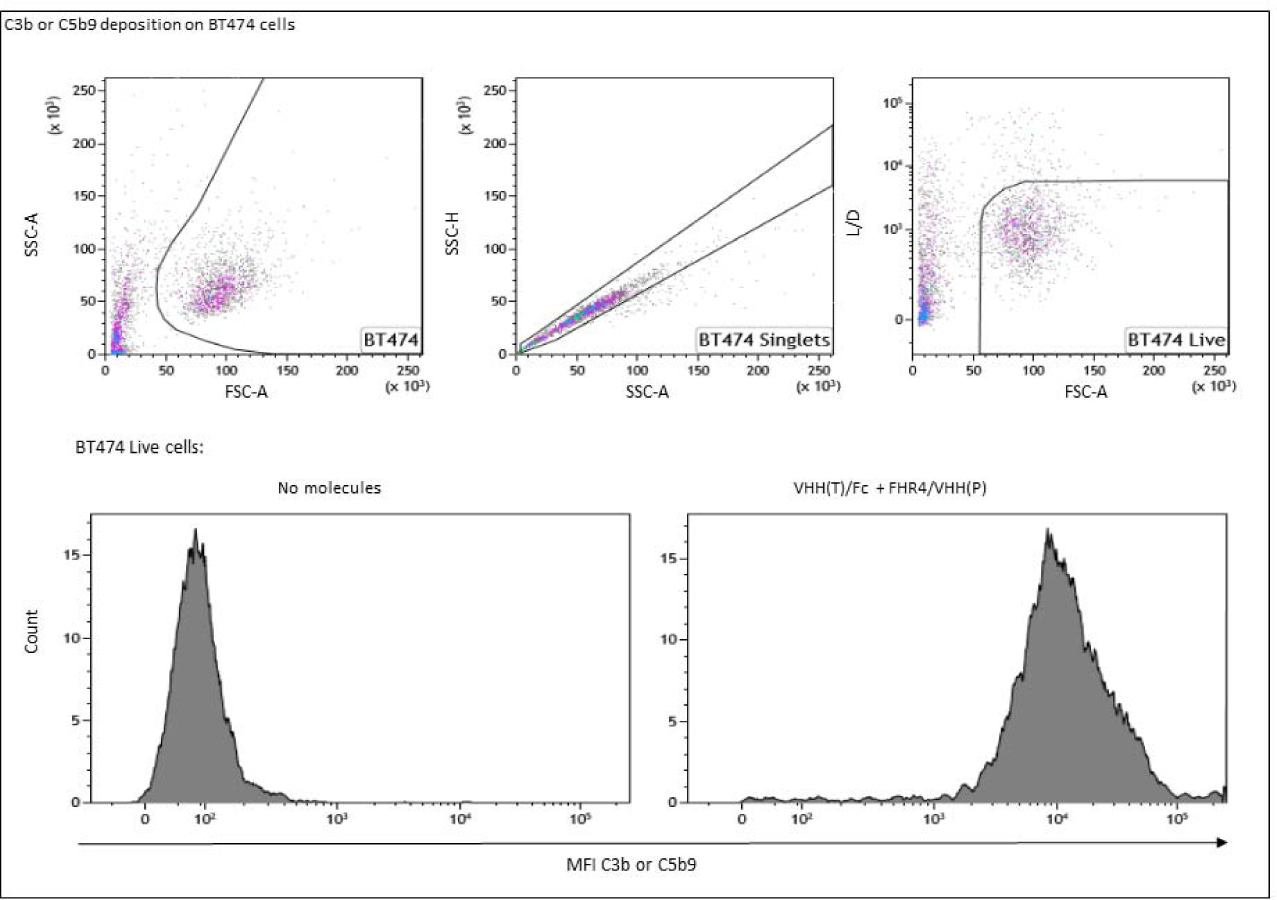
: Gating strategy for flow cytometry analysis of C3b, C5b9 depositions and Complement Dependent Cytotoxicity on BT474 cells. Live BT474 single cells were identified by an intersect gating strategy excluding debris (FSC-A/SSC-A), doublets (SSC-A/SSC-H) and dead cells (FSC-A/Live Dead). Histogram plots on live BT474 cells were used to represent C3b or C5b9 MFI. The percentage of dead cells was calculated by dividing the number of L/D-positive cells by the total number of cells analyzed.

**Supplementary figure 3.**
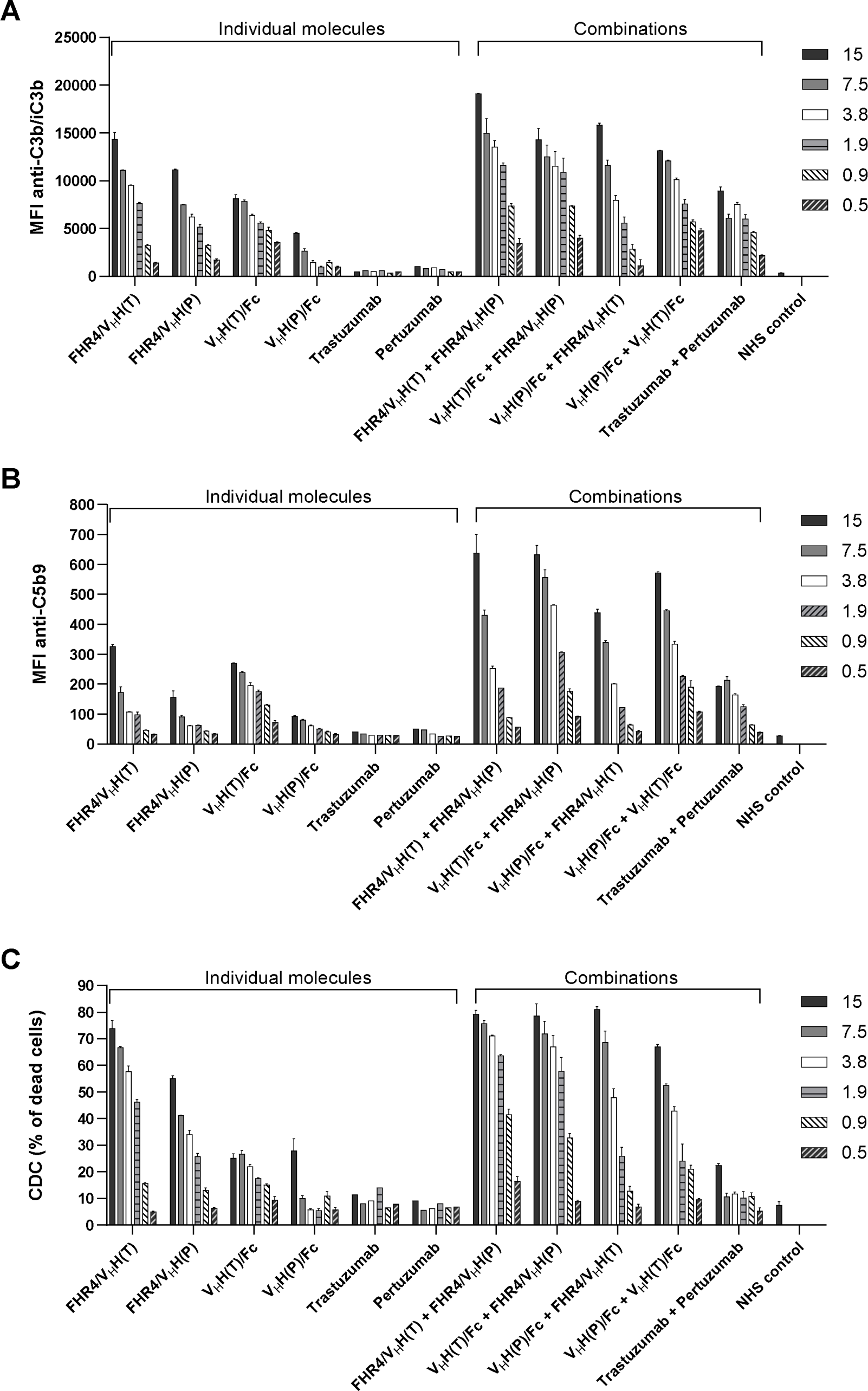
: Dose response analysis of C3b/iC3b deposition **(A),** MAC formation **(B),** and complement-dependent cytotoxicity **(C)** on BT474 tumor cells incubated with 3-fold serial dilutions: from 15 µg to 0.5 µg/well in case of individual molecules, and from 7.5 µg to 0.25 µg/well of each in case of molecule combinations. As controls, therapeutic antibodies and NHS were used. **(A)** C3b/iC3b deposition was detected with mouse anti-human C3b mAb and a secondary goat anti-mouse IgG Ab conjugated with AF647. CoMiX-Fc and CoMiX-FHR4 molecules elicit stronger complement activating effects than trastuzumab, pertuzumab and the combination of these two antibodies. Combining CoMiX-Fc and CoMiX-FHR4 molecules with other multimers resulted in the highest level of C3b desposition. **(B)** Staining with anti-C5b9 mAb followed by PE-conjugated anti-mouse IgG pAb was used to detect membrane attack complex (MAC) formation. **(C)** The percentage of dead cells was calculated by dividing the number of live/dead-positive (dead) cells with the total number of analysed cells. Data are presented as mean values ±SD of n = 3 independent experiments.

**Supplementary figure 4.**
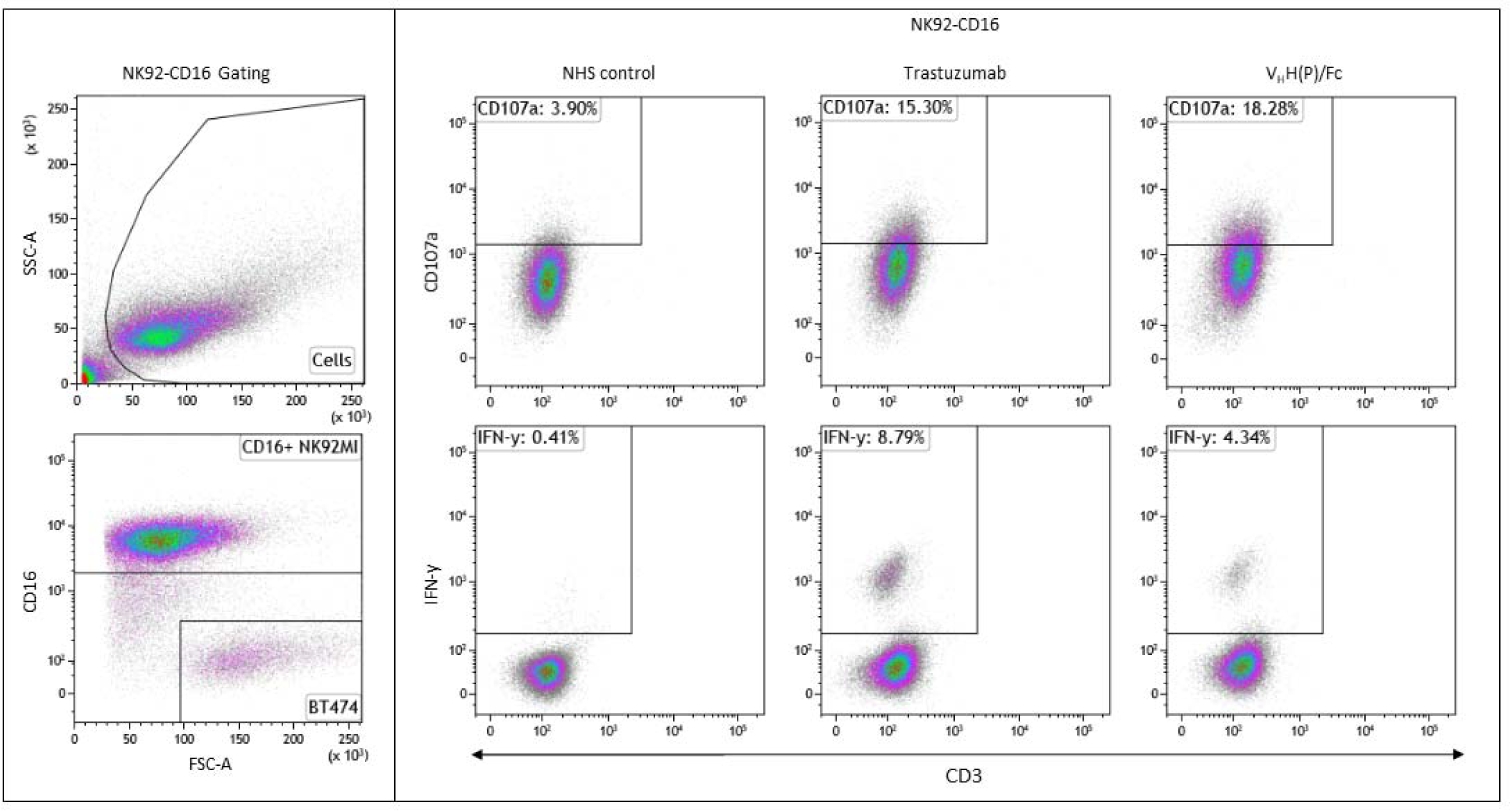
Flow cytometry dot plots representing the gating strategy and the expression of CD107a and IFN-y on live CD3-NK92^humCD16^ cells. Live NK92^humCD16^ cells were identified using an intersect gating strategy excluding debris (FSC-A/SSC-A), BT474 cells (FSC-A/CD16) and dead cells (FSC-A/Live Dead (data not shown). Representative dot plots on Live NK92^humCD16^ cells are depicted to show the percentages of CD107a or IFN-y expression on CD3-NK92^humCD16^ cells in presence of trastuzumab and V_H_H(P)/Fc.

## Supplemental material and methods

### Plasmid design

The pEF-IRESpac bicistronic expression vector^28^ was used to clone all expression cassettes into the multiple cloning site between EcoRI and XbaI to generate stable mammalian cell lines that express high levels of recombinant proteins and contains the puromycin resistant gene for selection of stably transfected cells. The expression cassettes were cloned as previously described^13^. Briefly, the signal peptide derived from the tumor necrosis factor receptor superfamily member 16 (UniProt no. P08138) was cloned between EcoRI and BglII, followed by the cDNA encoding the 2D3 V_H_H anti-HER2 (trastuzumab-competing epitope), 47D5 V_H_H anti-HER2 (pertuzumab-competing epitope), the FHR4A protein (UniProt no. Q92496) or eGFP, inserted between BglII and BspEI. The cDNA encoding (i) a 5× (SG4S) linker, followed by (ii) the C4bp C-terminal α-chain oligomerization domain and (iii) the Fc fragment from human IgG1 (hinge, CH2 & CH3), was cloned between BspEI and NotI. Plasmids containing the 2D3-V_H_H anti-HER2 (trastuzumab-competing epitope) and 47D5 (WO2009068625A2) V_H_H anti-HER2 (pertuzumab-competing epitope) adapted from patent WO2009068625A2, were synthesized by ProteoGenix SAS (Schiltigheim, France).

### Cell culture and antibodies

All CoMiX were produced in HEK293T cells (American Type Culture Collection, ATCC CRL-3216) cultured in Dulbecco’s modified Eagle’s medium (Gibco, Thermo Fisher Scientific) supplemented with 10% (v/v) heat-inactivated Fetal Bovine Serum (Gibco), 1 U/mL of penicillin (Westburg, Leusden, The Netherlands), 1 μg/mL of streptomycin (Westburg), and 4 mM of glutamine (Westburg). For protein production, Opti-MEM reduced serum media (Gibco) was used. Trastuzumab-sensitive BT474 (ATCC HTB-20), trastuzumab-resistant BT474 clone 5 (ATCC CRL-3247) cell lines were purchased from ATCC and NK92^humCD^^16^ provided by Pr Clémenceau, University of Nantes, France^29^ cells. Albumin Fraction V (BSA) was purchased from Roth (Keerbergen, Belgium). Goat anti-human IgG Fc pAb (#ab97221), goat anti-human IgG Fc HRP pAb (#ab97225), goat anti-rabbit IgG Fc AF568 pAb, and mouse anti-human C5b9 mAb (#ab66768, clone aE11) were purchased from Abcam (Cambridge, UK). Rabbit anti-His pAb was purchased from Bethyl (#A190214A, ImTec Diagnostic NV, Antwerpen, Belgium). Goat anti-mouse IgG AF647, the near IR Live/Dead staining, and DAPI were obtained from Invitrogen (Thermo Fisher Scientific BVBA, Merelbeke, Belgium). Mouse anti-human C3/C3b/iC3b mAb was provided by Cedarlane (#CL7637AP, Sanbio B.V., Uden, The Netherlands). Goat anti-human IgG AF647 was purchased from Southern Biotech (#2048-31, Birmingham, United States). GolgiStop and GolgiPlug were purchased from BD Biosciences (Eysins, Switzerland). The Brilliant Violet 421 anti-human CD107a and goat anti-mouse IgG AF488 antibodies were purchased from Biolegend (Amsterdam, The Netherlands). Rabbit anti-human C3d polyclonal antibody was provided by Agilent (DAKO, #A0063, Amstelveen, The Netherlands). Mouse anti-human FHR4 AF488 mAb was purchased from R&D Systems Europe Ltd, (#IC5980G-100UG, clone 640212, Bio-Techne, Abingdon, UK). Goat anti-mouse IgG APC was obtained from Jackson ImmunoResearch (Sanbio). Commercial trastuzumab (Herceptin) and pertuzumab (Perjeta) antibodies were purchased from Roche (Prophac, Howald, Luxembourg).

### Protein production and purification

After selection of the clones, clones were produced during 3 cycles of 48h in serum-free optiMEM followed by 24h in complete DMEM medium. Opti-MEM supernatants were precleaned by centrifugation (4000 rpm/10 min.) and filtered using 0.22 µm PVDF vacuum filter units (GE-Healthcare, Leuven, Belgium). CoMiX-Fc molecules were purified using Protein G Sepharose® 4 Fast Flow beads (GE Healthcare). Filtered supernatants were incubated with Protein G beads roller-bottled for 2 days at 4°C. Beads were collected by centrifugation and washed 2 times with 50 ml PBS. CoMiX-Fc were eluted using phosphate citrate buffer at a low pH (pH 2.7). The eluate was immediately neutralized with buffer (bi-carbonate buffer, pH 9.1) at a 4:1 ratio to a final pH 7.2-7.4. Protein solutions were next concentrated using Amicon© 30 kDa MWCO centrifugal filters (Millipore-Merck Chemicals NV/SA). The protein concentration was then measured at 280 nm on a NanoDrop™ micro-volume spectrophotometer.

CoMiX-FHR4 molecules and V_H_H controls were purified using Nickel His-Trap™ Excel columns (Cytiva). Filtered Opti-MEM supernatants were supplemented with imidazole (Sigma-Aldrich) to a 20 mM final concentration to avoid nonspecific binding. The molecules were then loaded to the His-Trap column for 3 days using a peristaltic pump with a flow rate of 1-2 ml/min. (GE-Healthcare, VWR). The His-Trap column was connected to a NGC Chromatography purification system (Bio-Rad) and washed with 20 mM phosphate buffer with 500 mM NaCl at pH 7.2 supplemented with imidazole for elution. Three elution fractions (f1-f3) were collected after purification and analyzed to maximize the FHR4-valency as high valencies are a prerequisite to overcome complement inhibition threshold^13^. Fraction 1 (f1) was collected under 120 mM imidazole concentration, whereas for f2 and f3 1 M imidazole concentration was used. As the molecular pattern of f2 and f3 was identical (supplementary figure 1A), the two fractions were pooled and used for further experiments. The elution fraction was concentrated on Amicon© 30 kDa MWCO centrifugal filters and dialyzed in 3×5L of PBS using 10 kDa MWCO Slide-A-lyzer® dialysis cassettes (Thermo Fisher Scientific) to remove any residual imidazole. Protein concentration was measured on Nanodrop (Ozyme, Saint-Cyr-l’École, France).

### SYPRO Ruby protein gel staining and western blotting

To visualize the molecular pattern of CoMiX molecules, we used SDS/PAGE followed by either Western blotting or SYPRO Ruby gel staining. One µg of purified CoMiX molecules were mixed with 4 µl of 4X Laemmli sample buffer (Bio-Rad), supplemented with 10% 2-mercaptoethanol (Sigma-Aldrich) for reducing conditions. The samples were loaded onto 4-15% Mini-Protean® TGX™ protein acrylamide gels (Bio-Rad) and electrophoresed. After fixation (methanol/acetic acid), the gel was stained using SYPRO Ruby staining solution according to the manufacturer’s instructions. The gel was then revealed by the Amersham™ Typhoon™ biomolecular imager with appropriate filter. For western blot analysis, the proteins were electroblotted from gels onto methanol-activated Amersham Hybond 0.2 μm low-fluorescence background PVDF membranes (GE Healthcare) using a Mini Trans-Blot® electrophoretic transfer cell system (Bio-Rad). The membrane was blocked with 5% (w/v) low-fat milk in PBS overnight at 4°C shaking. The next day, 1 µg of the following antibodies were added: mouse anti-human FHR4 AF488-conjugated monoclonal Ab (mAb) (R&D Systems) for CoMiX-FHR4 molecules and goat anti-human IgG polyclonal antibody (pAb) AF647-conjugated (Abcam) for CoMiX-Fc molecules. After 3 washes with 0.05% TBS-Tween buffer, the membranes were dried and analysed using the Amersham™ Typhoon™ laser-scanner platform.

